# CRISPR-Cas Inhibits Natural Transformation Through Altruistic Group Defense and Self-Sacrifice

**DOI:** 10.1101/2021.09.16.460680

**Authors:** Robert M. Cooper, Jeff Hasty

## Abstract

CRISPR-Cas systems present an evolutionary tradeoff: does defense against phages and other parasitic DNA also prevent cells from acquiring potentially helpful new genes? Genomic analyses of this conundrum have arrived at often contradictory conclusions. Meanwhile, experimental studies have focused mainly on phages, conjugation, or artificial transformation, but less work has examined natural competence, a major driver of evolution and antibiotic resistance. Here, we use *Acinetobacter baylyi*, which combines high natural competence with a functional CRISPR-Cas system, to experimentally probe the interactions between CRISPR-Cas and natural competence. In these bacteria, the endogenous CRISPR array largely allows natural transformation by targeted DNA. However, CRISPR-Cas then kills the newly autoimmune cells in a form of programmed cell death. CRISPR-Cas often allows self-targeting cells to form colonies, albeit with fitness costs. Thus CRISPR-Cas appears to block natural transformation in a process more akin to altruistic group defense than an individual immune system.

## Introduction

CRISPR-Cas systems are generally described as adaptive immune systems for prokaryotes^1^. CRISPR arrays store a series of short DNA target sequences, or spacers, which often derive from invading phage or mobile genetic elements. CRISPR-associated Cas proteins search for DNA matching these stored sequences, which they often cleave or shred upon detection. In contrast to CRISPR-Cas, which blocks foreign DNA, natural competence, another widespread phenomenon among bacteria, actively brings foreign DNA inside the cell^2^. If the foreign DNA integrates into the genome or re-circularizes into a replicating plasmid, the cell acquires potentially beneficial new genes. Thus, CRISPR-Cas systems and natural competence exert opposing effects on horizontal gene transfer (HGT): one blocking HGT, and the other causing it.

The apparently contrasting effects of CRISPR-Cas and natural competence present an evolutionary conundrum^3–8^. How do bacteria balance the competing imperatives of rapid adaptation on the one hand, versus protection against parasitic DNA on the other? Genomics-based evolutionary analyses have come to often contradictory conclusions. In some analyses, CRISPR-Cas does appear to slow adaptation via HGT, particularly when it comes to genes for virulence^9^ and multi-drug resistance^10,11^. However, in other analyses CRISPR-Cas did not reduce HGT over evolutionary time^12,13^. Even within a single study, the relationship between CRISPR-Cas and HGT can be complex; in *Acinetobacter baumannii*, CRISPR-Cas is simultaneously associated with both fewer plasmids, as expected, but also more genes related to biofilm formation^14^.

When it comes to natural competence in particular, the evolutionary relationship between CRISPR-Cas and HGT is especially muddled. As for HGT in general, natural competence seems to have a contradictory or context-specific relationship to CRISPR-Cas in evolutionary analyses^8^. CRISPR-Cas is prevalent among most Streptococci, but is conspicuously absent from highly competent *Streptococcus pneumoniae*, suggesting CRISPR-Cas is detrimental for species evolved for natural competence^15^. However, *Neisseria meningitidis* and *Aggregatibacter actinomycetemcomitans* exhibit the opposite: a positive correlation between CRISPR-Cas and natural competence^16,17^.

Contributing to the uncertain ecological relationship between CRIPSR-Cas and natural competence is the limited number of experimental studies. Much experimental work has focused on CRISPR-Cas defending against phage, conjugating plasmids, or laboratory electroporation, but natural competence, one of the three major routes of HGT, has received far less attention^15,16^. Here, we experimentally probe the relationship between CRISPR-Cas and natural competence in *Acinetobacter baylyi*, which combines a functional Type I-F CRISPR-Cas system^18–20^ with exceptional natural competence^21^. In this scenario, the result of a showdown between CRISPR-Cas and natural transformation turns out not to be as simple as a count of transformed cells, and the ecological function of CRISPR-Cas is not as clear-cut as immunity or self-defense.

## Results

### Most of endogenous CRISPR array does not block transformation

*A. baylyi* strain ADP1 contains a natural, 90-spacer CRISPR array^18^. To test the ability of the endogenous array to block natural transformation, donor DNAs were constructed containing one of 9 targeted protospacers from across the array, a random protospacer-length sequence, or no insert, placed immediately upstream of an antibiotic resistance marker. One set of donor DNAs had homology arms directing them to a neutral site on the genome, and another set had homology arms flanking the endogenous CRISPR array, so that the array would be deleted upon recombination with the donor DNA. These donor DNA constructs were incubated with both wild-type and Δ*CRISPR* cells, in which the CRISPR array had been deleted, and transformants were counted (Supplemental Figure S1). Transformation rates were then normalized to donor DNA with no protospacer (Figure 1).

**Figure 1:**
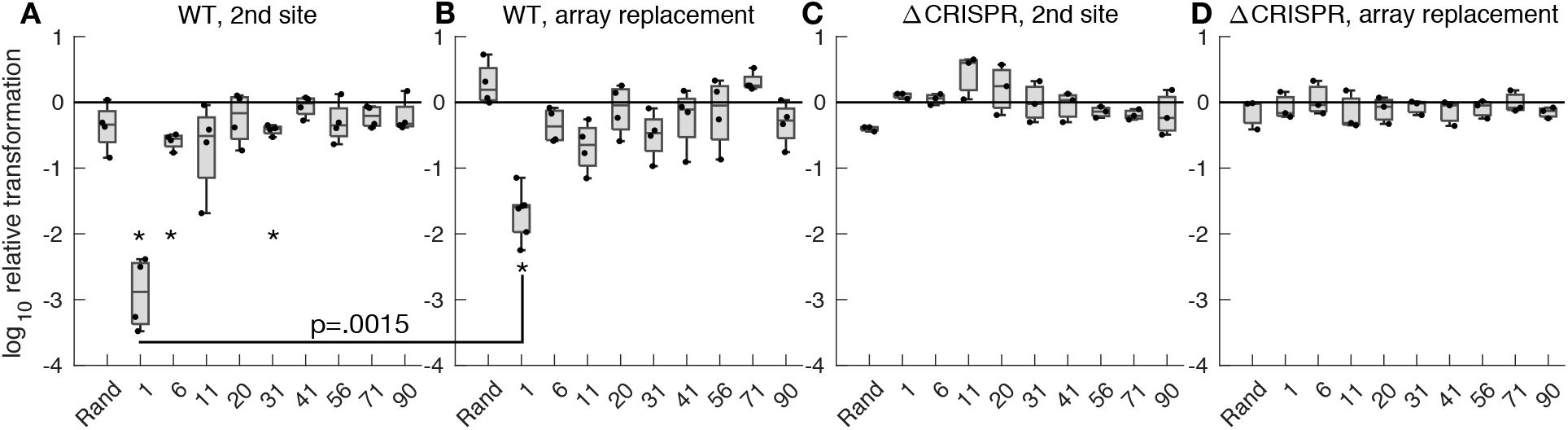
Relative transformation by DNA targeted by the endogenous CRISPR array. Transformation rates (transformants per recipient) for donor DNA containing various targets matching spacers from the endogenous array, or a random spacer, were normalized against transformation rates for donor DNA without a target. Donor DNA either inserted into a separate genomic site (A, C), or replaced the CRISPR array upon insertion (B, D). Recipient cells were either wild type (A, B) or controls lacking the CRISPR array (C, D). *: p<.00125 (Bonferroni corrected for 40 tests), one-tailed t-tests. Note the data is log scale. See also Figure S1.

After a Bonferroni correction for multiple hypothesis testing, CRISPR-Cas only significantly reduced transformation into a second locus for spacers 1, 6 and 31 (Fig. 1A). When the donor DNA replaced the CRISPR array, only spacer 1 allowed CRISPR-Cas to significantly reduce transformation (Fig. 1B). No donor DNAs had reduced transformation in control Δ*CRISPR* cells (Fig. 1C,D). Sequencing of genomic DNA from transformants showed that the CRISPR target and PAM sites were intact, except for those targeted by spacer 1, for which the target site was mutated.

This data showed that the endogenous CRISPR array is functional – the first spacer reduced natural transformation by targeted DNA by roughly 1000-fold – but the efficacy dropped by 2 orders of magnitude by just the 6^th^ spacer in the array, rendering most of the array at best minimally effective at preventing transformation. Interestingly, the highly effective spacer 1 was 20-fold less effective against incoming DNA that deleted the CRISPR array upon recombination (Fig. 1A vs 1B). This suggested that CRISR-Cas was acting largely after recombination in to the genome had occurred. Once incoming DNA had replaced the CRISPR array, production of crRNAs would cease, and the inserted DNA would need only to avoid CRISPR-Cas targeting for a few cell divisions, after which existing crRNAs would be diluted by growth.

### Deleting CRISPR spacers helps foreign DNA avoid targeting

To test this hypothesis with more CRISPR spacers, we placed single CRISPR spacers targeting a kanamycin resistance gene, a random spacer, or an array containing all 4 targeting spacers into a separate neutral locus of the *A. baylyi* genome. These strains were then incubated with donor DNA containing the targeted resistance gene flanked by homology arms to either a second site in the genome (Fig. 2A, Supplemental Fig. S2A) or to the same site as the introduced array (Fig. 2B, Supplemental Fig. S2B), and transformants were counted and normalized to cells containing a random spacer. One spacer (B1) did not significantly reduce transformation relative to a random spacer, but the other three spacers and the 4-spacer array did. Two spacers (T1 and T2), were 10-fold less effective against donor DNA that replaced the array, as predicted if CRISPR acts after integration.

**Figure 2:**
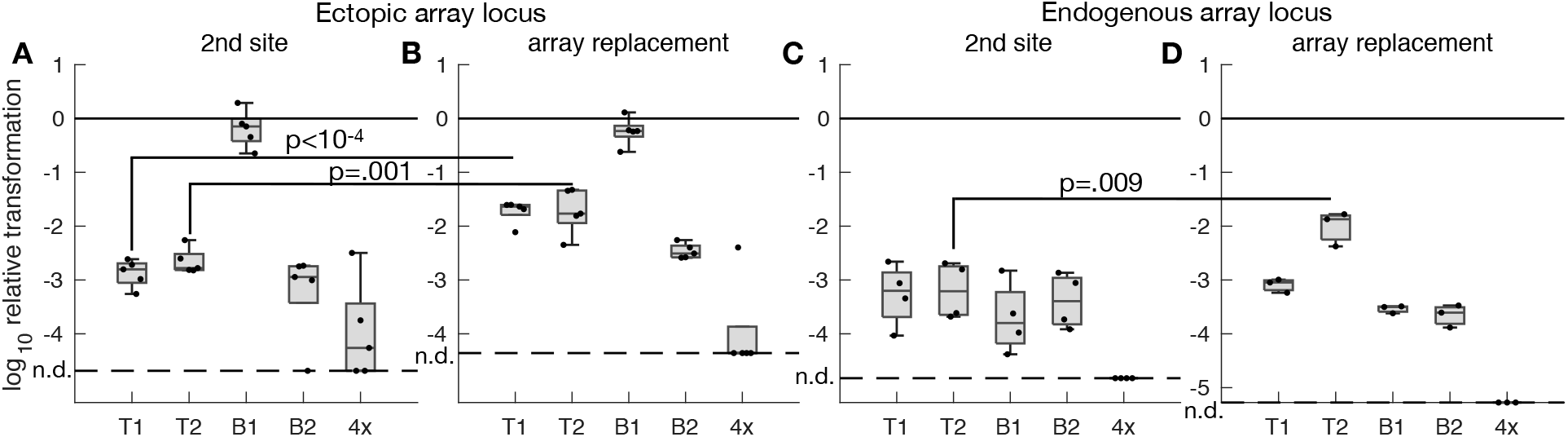
Relative transformation by DNA targeted by introduced CRISPR spacers. Transformation rates (transformants per recipient) were normalized against rates for cells containing an introduced random spacer. CRISPR spacers were introduced into an ectopic locus (A, B) or used to replace the endogenous locus (C, D). Donor DNA either inserted into a separate genomic site (A, C) or replaced the introduced spacers (B, D). P-values from two-sample, one-tailed t-tests. Dashed lines indicate the limit of detection, and data below that limit is shown as n.d. for none detected. Note the data is log scale. See also Figure S2.

Multiple crRNAs in the same cell compete for the Cas proteins, thus competitively inhibiting each other^22^. Therefore, to provide a best-case scenario for the CRISPR-Cas system, we replaced the entire CRISPR array with the same single spacers or 4-spacer array targeting the kanamycin resistance gene (Fig. 2C,D, Supplemental Fig. S2C,D). The single spacers reduced transformation by roughly 1000-fold, similar to the single spacers in an ectopic locus or the first spacer in the natural array, and a 4-spacer array was more effective than any single spacer. Spacer T2 was again less effective against array-replacing DNA than against DNA inserting into a second site, although this time spacer T1 was not. Thus, even in the absence of competitive inhibition from a long array, the *A. baylyi* CRISPR-Cas system shows evidence of acting after genomic integration.

### CRISPR “escapes” suffer fitness costs

Much of the endogenous CRISPR array did not significantly reduce natural transformation by targeted DNA, as measured by transformed colony-forming units (CFUs). At first glance, this would seem to suggest the 90-spacer array is mostly useless. However, even though the transformed cells can grow into colonies, the CRISPR-Cas system could still reduce the prevalence of this DNA in the population over time by killing some transformed cells, which would lower their effective growth rate. To test this, we performed a pulse-chase experiment, exposing *A. baylyi* cells to a 30-minute pulse of donor DNA containing a single protospacer targeted by the endogenous array, and counting the proportion of transformed cells both immediately and after 4 hours of exponential growth.

As before, the CRISPR target sequences did not significantly reduce transformation immediately after the 30-minute DNA pulse, except for protospacers 1 and 6 (Fig. 3A, Supplementary Fig. 3). However, spacers throughout the first half of the array did reduce fitness following transformation with their targets (Fig. 3B). After a Bonferroni correction, spacers 6, 11, 20, 31, and 41 significantly reduced fitness relative to transformants with no protospacer. The relative fitness of transformants targeted by spacer 1 could not be calculated, since none were detected at the first time point. The three spacers from the second half of the array (56, 71, and 90) had no fitness effect on self-targeting transformants during exponential growth.

**Figure 3:**
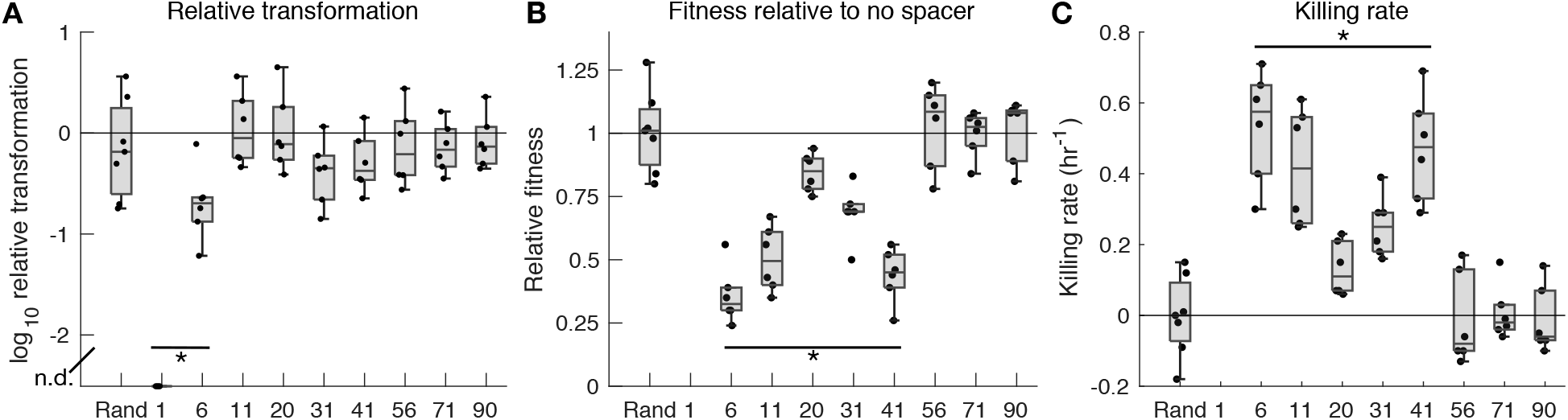
Fitness cost in CRISPR “escape” transformants. A) Transformation rates (transformed CFUs per recipient cell, log scale) for donor DNA containing indicated CRISPR targets (x-axis) at the first time point of pulse-chase experiments. Data below detection is shown at the limit of detection, denoted n.d for none detected. B) Relative fitness of transformants, with respect to cells transformed by donor DNA without a protospacer (1 indicates equal fitness). C) CRISPR-Cas killing rates for transformants containing indicated targets. Note that relative fitness (B) and killing rate (C) could not be calculated for spacer 1. *: p<.005, one-sample, one-tailed t-tests. See also Figure S3.

Under the assumption that the fitness cost is due to the CRISPR-Cas system killing transformed cells, we can also calculate the killing rate. All spacers in the first half of the CRISPR array killed cells containing their targets at rates significantly greater than 0 (Fig. 3C). The killing rate for spacer 1 could not be calculated, but it was presumably greater than the average growth rate of transformed cells with no spacer, which was 0.76 hr^−1^.

### CRISPR-Cas defense causes genomic DNA damage

If the reduced fitness of transformants containing CRISPR-targeted DNA results from ongoing genomic self-targeting, those cells should suffer extensive genomic DNA damage. To test this, we inserted a GFP gene driven by the *recA* promoter into the *A. baylyi* genome as a reporter for DNA damage^23^. To maximize the signal, we engineered recipient cells so that the only transferring region was the 32 bp protospacer, since small inserts integrate much more efficiently than larger ones. We mixed cells with DNA, spotted them on agar, and observed them using time-lapse microscopy through a glass-bottom dish. To confirm high transformation efficiency in our setup using a small insert, we first visualized repair of a constitutively expressed, but broken, GFP gene, using donor DNA that repaired 3 nearby stop codons in GFP (note this demonstration construct is separate from the GFP DNA damage reporter). Under the microscope, a visibly large proportion of cells were transformed (Supplemental Fig. S4).

Having confirmed high transformation efficiency in these experimental conditions, we returned to the strain containing the *P_recA_*-*GFP* reporter for DNA damage. A small fraction of cells expressed GFP even when incubated without DNA or with DNA without a CRISPR target (Fig. 4A). Unlike a previous study, in which uptake of long genomic DNA induced signīficant genome damage^23^, short 3 kb fragments did not increase DNA damage when they lacked a CRISPR target. However, when cells were incubated with donor DNA containing a target from the CRISPR array, far more cells activated the DNA damage response pathway (Fig. 4B-E, Supplementary Figures S6-S14). Interestingly, when the cells were transformed with the target for spacer 1, the DNA damage was generally confined to single cells (Fig. 4B). However, as the target in the donor DNA came from farther back in the CRISPR array, the DNA-damaged cells appeared in larger clumps (Fig. 4D,E). In time lapse movies, these clumps appeared to begin expressing GFP around the same time, generally when the local growth rate of the expanding microcolony slowed down (Supplemental Movies).

**Figure 4:**
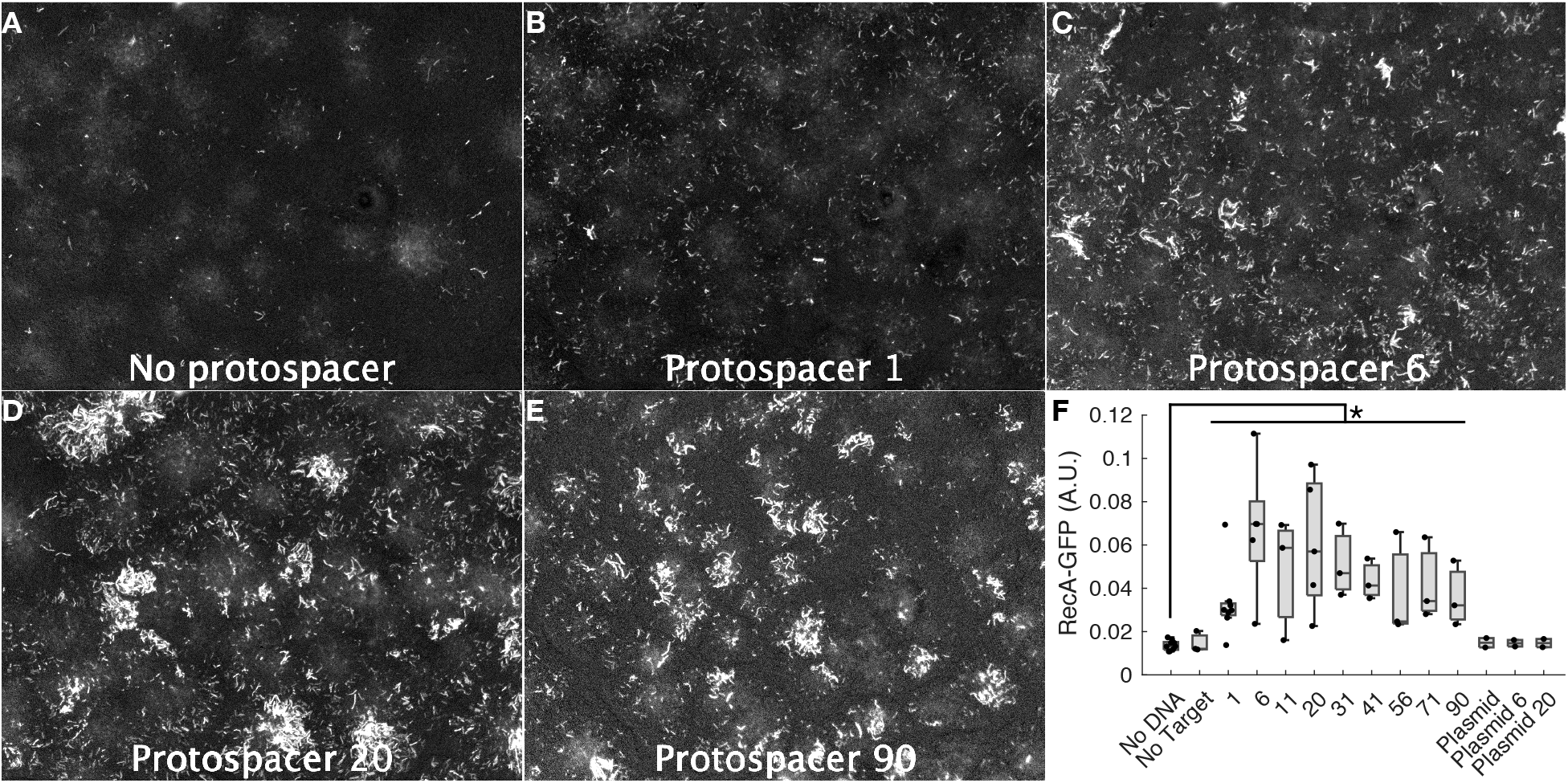
Natural transformation of CRISPR-targeted DNA causes genomic damage. GFP reporter cells for DNA damage were incubated under agar with donor DNA with no CRISPR targets (A), or targets matching spacer 1 (B), 6 (C), 20 (D), or 90 (E) from the endogenous CRISPR array. F) Fraction of cell-occupied pixels that were GFP-positive (out of 1), indicating DNA damage, when *P_recA_*-*GFP* reporter cells were incubated alone; with genomically integrating donor DNA containing no target or targets from the endogenous array; or with non-integrating plasmid DNA containing no target or targets from the endogenous array. *: p<.005, two-sample, one-tailed t-test. See also Figures S4-S23.

To quantify DNA damage, we counted the fraction of cell-occupied pixels that were above a threshold brightness in the GFP channel (Fig. S5, see Methods). Experiments without DNA or with donor DNA without a CRISPR target had very few positive pixels, but when donor DNA contained a CRISPR target, there were many more positive pixels, although with significant variation across experiments (Fig. 4F). All targeted protospacers caused significantly more DNA damage than experiments without donor DNA.

In principle, the observed DNA damage response could be due to the CRISPR-Cas system targeting incoming DNA either before or after the DNA integrated into the genome. To exclude targeting of DNA outside the genome, we also used a plasmid as donor DNA, either without a CRISPR target or with protospacers 6 or 20. These protospacers had generated significant DNA damage signal for integrating DNA, but in a non-integrating plasmid of similar length, they did not cause significant DNA damage (Fig. 4F). Since CRISPR-Cas targeting of a 3 kb DNA fragment in the cytoplasm did not activate the RecA response, the DNA damage signal for integrating DNA most likely came from genomic self-targeting following DNA integration.

## Discussion

The widely used metaphor of an adaptive immune system implies that CRISPR-Cas defends the individual cell against parasitic DNA. In 2013, Johnston *et al*. proposed a model in which CRISPR-Cas antagonizes natural competence not by preventing transformation, but by killing transformed cells, in contrast to the immunity metaphor^24^. While this paper has received several citations, only 3 mentioned the hypothesis of CRISPR-Cas acting after transformation, killing cells rather than defending them^25–27^. Previous work has shown that CRISPR-Cas can kill cells through genomic self-targeting, but not in the context of natural transformation by exogenous DNA targeted by an endogenous CRISPR-Cas system^28–32^. Here, we present several lines of experimental evidence that in *A. baylyi*, much of CRISPR-Cas interference with natural transformation does indeed occur after the transformation event.

In all CRISPR arrangements tested here – the endogenous array (Fig. 1), ectopic spacers (Fig. 2A,B), or introduced spacers in place of the endogenous array (Fig. 2C,D) – at least some spacers were less effective at blocking same-site, array-replacing donor DNA than second-site insertions, as expected if CRISPR acts after integration. However, this was not the case for all spacer-target pairs. For some spacers, CRISPR-Cas interference was too weak to reduce transformation much at either site. For others, CRISPR-Cas interference may have been strong enough that even for array-replacing insertions, mutations were the main source of transformants. Such variation in spacer effectiveness seen here is expected given previous results. Early spacers in an array are expressed more strongly than later ones, and thus work more effectively^33^. Also, local sequence can create context-dependent variation in pre-crRNA processing^34^, which may explain why different spacers were more effective when expressed from different genomic loci (Fig. 2A,B vs Fig. C,D).

The data here does not exclude the possibility that some incoming DNA could be blocked by CRISPR-Cas before genomic integration. However, significant genomic damage was observed for both the first and last endogenous spacer in the *P_recA_*-*GFP* experiments (Fig. 4), suggesting that self-sacrifice is always a feature of the CRISPR-Cas system in *A. baylyi*. Spacer 1 caused less GFP signal than other spacers from the first half of the array, but this does not necessarily mean that it was more effective at preventing initial genomic integration. For many weaker spacers, DNA damage signal occurred in clumps, which likely originated from a single transformation event whose host cell was able to divide several times before CRISPR-Cas killed its descendants, generating GFP signal from many cells for one transformation. For this reason, P*_recA_*-GFP signal is not linearly related to spacer strength.

CRISPR-Cas search kinetics support this interpretation. In *E. coli*, the average search time for a single Cas9 molecule or a Type I Cascade complex to find its target has been estimated at 1.5 and 6 hours, respectively^22,35^. The search time can be lowered in practice if multiple CRISPR-Cas complexes are loaded with the same guide RNA and search in parallel, but the kinetic data supports our observations that self-targeting transformants can escape death for several divisions during more rapid growth. This is similar to the case for plasmids, whose replication is better able to outpace CRISPR-Cas cleavage during rapid growth^36^.

These results argue that one must use caution when measuring CRISPR-Cas effectiveness against natural competence. Most endogenous spacers had no effect on natural transformation when counting colony-forming units (CFUs). However, closer examination of growth rates or genome damage revealed that CRISPR-Cas was indeed working to steadily reduce the population prevalence of targeted transformants. This observed post-integration action by CRISPR-Cas demonstrates a limitation of the immune system metaphor. For both the strongest and the weakest spacers, CRISPR-Cas often kills cells rather than defending them, acting more like a self-destruct switch, or abortive infection. This is often true for CRISPR-Cas defense against phages as well^37,38^. This distinction is important beyond accurate semantics, since altruistic group defense produces more complex ecological dynamics, and is subject to more complex selective forces, than pure self-defense^39^.

## Acknowledgements

This work was supported by NIH grant R01GM069811.

## Author contributions

Conceptualization, R.C., J.H.; Methodology, R.C; Investigation, R.C.; Writing – Original Draft, R.C.; Writing – Review & Editing, R.C., J.H; Visualization, R.C.

## Methods

### Cell culture and cloning

*A. baylyi* strain ADP1 was obtained from the American Type Culture Collection (ATCC #33305). Cells were propagated in standard LB media at 30 or 37 °C. DNA constructs were prepared using Gibson Assembly or Golden Gate cloning methods to generate linear DNA, which was used to transfom *A. baylyi* directly, by incubating 20-100 μl cells plus DNA for 30 minutes to a few hours. Clones were picked from selective plates, genomic DNA was purified using a genomic DNA miniprep kit, regions of interest were amplified by PCR, and the transformants were verified by Sanger sequencing. For *P_recA_*-*GFP*, clones were screened for low-level GFP fluorescence by placing the plate on a blue transilluminator, rather than antibiotic selection. Multiplex CRISPR arrays were assembled using a method described previously ^19^.

Three genomic loci were used in the experiments described here. One was the natural CRISPR array located between ACIAD2484 and ACIAD2500. A second was the gene remnant ACIAD2826, which was replaced as a neutral locus for insertions, denoted *ntrl1*. A third was a neutral locus used previously, replacing ACIAD2186, 2187, and part of 2185, into which the ectopic CRISPR arrays were inserted^40^. For ectopic CRISPR arrays, the inserts included 684 bp upstream of the first repeat from the endogenous array. For *P_recA_*-*GFP*, the insert included 214 bp of promoter from upstream of the *recA* gene, as described previously^23^.

### Transformation experiments

*A. baylyi* was grown overnight in LB broth at 30°C with shaking. Just before experiments, cultures were washed and resuspended in fresh LB. 20 μl of cells were mixed with 40 ng of purified DNA and incubated at 30°C with shaking for 2 hours. Samples were then serially 10-fold diluted and spotted on LB agar plates with and without selective antibiotics (35 μg/ml kanamycin or spectinomycin), with 3 technical replicates for each dilution level. Plates were incubated at 30°C overnight, and colonies were counted at the lowest dilution level that still had clearly separated colonies. Counts for the three technical replicate spots were averaged. Each experiment was repeated at least 3 times, on at least 2 separate days.

For pulse-chase experiments, cells were growth overnight at 30°C, washed and diluted 1/13x into fresh LB, and re-grown for 3 hours at 37°C. The cells were then washed into fresh LB again, divided into 20 μl aliquots, mixed with 40 ng DNA, and incubated at 37°C. After 30 minutes, DNase and DNase buffer were added, and incubation continued for another 15 minutes at 37°C to digest remaining donor DNA. The cultures were then diluted into 600 μl fresh LB, and incubated at 37°C, with aliquots taken for serial dilution and spotting after 15 minutes and 3-4 hours.

Relative fitness was calculated using the equation 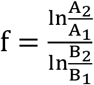, where *A* is cells transformed by DNA containing a given target, *B* is cells transformed by control DNA without a protospacer, and the subscripts indicate time points. Killing rates were calculated as 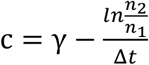, where *γ* is the growth rate of transformants with non-targeted control DNA without a protospacer, and *n* is the number of transformants at each time point.

### Statistics

Serial dilutions produce data best compared and plotted on a log scale, so data were plotted and compared after taking base 10 logarithms. Hypothesis testing was performed in Matlab using single-tailed t-tests. Where multiple samples were tested at once, the standard significance threshold of p<.05 was divided by the number of samples (Bonferroni correction for multiple hypothesis testing). Where data points were below detection, they were manually set to the limit of detection as the most conservative way to include them in the statistical analysis.

### Visualizing DNA damage

The construct *P_recA_*-*GFP* was prepared containing 214 bp of the *A. baylyi recA* promoter fused to GFP^23^, with approximately 1 kb homology arms surrounding a *vgrG* operon, and transformed into *A. baylyi.* This is a Type VI secretion system toxin-antitoxin operon, so in the absence of cells with the toxin, this insertion is neutral. In addition, cells contained a spectinomycin resistance gene in another neutral locus. Donor DNA for visualization experiments consisted of 3 kb covering that spectinomycin resistance gene and surrounding DNA, with a 32 bp protospacer inserted before the spectinomycin gene. Thus, the donor DNA had a 32 bp insert embedded within 3 kb of total homology to recipient cells. Cells were grown overnight in LB with shaking at 30 °C, washed, diluted 1:20 into fresh LB, and mixed 1:1 with donor DNA. Agar pads were prepared by pouring molten LB containing 2% wt/vol agar into 35 mm glass-bottom dishes, allowed to solidify, scooped out with a spatula, and placed upside-down. Spots of 0.5 μl of the cell-DNA mixtures were deposited onto the raised pads created from the glass window cutout and allowed to dry, after which the pad was placed back into the glass-bottom dish. Each experimental run contained 6 spots, including a no-DNA control. Three experiments were run for each CRISPR target and two for each plasmid. An additional 5 spots were imaged for protospacer 1 to obtain higher statistical power. A total of 9 no-DNA control spots were imaged, one for each experimental run. Experiments using plasmid DNA also included integrating DNA spots as positive controls. Time-lapse movies were obtained on a Nikon TE microscope using a 20x objective, with a 30-minute time step. Time points for analysis were selected at the point when growth significantly slowed, generally after 36 hours. GFP images were analyzed using the following procedure:

- Preprocessing: Images were preprocessed by dividing by a blank background image, which was itself first smoothed with a Gaussian filter with a radius of 5 pixels.
- Excluding empty areas: Regions with no cells were outlined manually, refined using the Image Segmenter app in Matlab, and excluded from further analysis.
- Background autofluorescence: To compensate for increased background fluorescence in the centers of microcolonies, an image-specific background was constructed. First, a Gaussian curve was constructed using the mean and variance of the GFP pixel histogram. Pixels greater than the 99th percentile of that curve were suppressed down to the 99th percentile level. The resulting image was smoothed with a Gaussian filter of radius 20 pixels to create an autofluorescence background. This was subtracted from the pre-processed GFP image, yielding the final background-corrected image.
- Histogram normalization: A histogram of GFP pixel intensity was constructed, and a new Gaussian curve was fit to the lower 60% of pixels, representing an expected distribution in the absence of GFP expression. The histogram was shifted laterally to be centered at 0, its width was normalized to σ = 1, and its height was normalized such that the total area under the curve was 1, to allow visual histogram overlays (Supplemental Fig. S5).
- Bright pixels: Bright pixels were defined as those greater than the 99.5 percentile of the aligned Gaussian.

## References

1. Makarova, K. S. et al. An updated evolutionary classification of CRISPR-Cas systems. Nature Reviews Microbiology 13, 722–736 (2015).

2. Johnsborg, O., Eldholm, V. & Håvarstein, L. S. Natural genetic transformation: prevalence, mechanisms and function. Res Microbiol 158, 767–778 (2007).

3. Weinberger, A. D. & Gilmore, M. S. CRISPR-Cas: To Take Up DNA or Not—That Is the Question. Cell Host & Microbe 12, 125–126 (2012).

4. Westra, E. R. et al. The CRISPRs, they are a-changin’: how prokaryotes generate adaptive immunity. Annual review of genetics 46, 311–339 (2012).

5. García-Martínez, J., Maldonado, R. D., Guzmán, N. M. & Mojica, F. J. M. The CRISPR conundrum: evolve and maybe die, or survive and risk stagnation. Microbial Cell 5, 262–268 (2018).

6. Jiang, W. et al. Dealing with the Evolutionary Downside of CRISPR Immunity: Bacteria and Beneficial Plasmids. PLOS Genetics 9, e1003844 (2013).

7. Hatoum-Aslan, A. & Marraffini, L. A. Impact of CRISPR immunity on the emergence and virulence of bacterial pathogens. Current Opinion in Microbiology 17, 82–90 (2014).

8. Bondy-Denomy, J. & Davidson, A. R. To acquire or resist: the complex biological effects of CRISPR–Cas systems. Trends in microbiology 22, 218–225 (2014).

9. García-Gutiérrez, E., Almendros, C., Mojica, F. J. M., Guzmán, N. M. & García-Martínez, J. CRISPR Content Correlates with the Pathogenic Potential of Escherichia coli. PLoS ONE 10, e0131935 (2015).

10. Aydin, S. et al. Presence of Type I-F CRISPR/Cas systems is associated with antimicrobial susceptibility in Escherichia coli. Journal of Antimicrobial Chemotherapy 72, 2213–2218 (2017).

11. Palmer, K. L., Gilmore, M. S. & Losick, R. Multidrug-Resistant Enterococci Lack CRISPR-cas. mBio 1, e00227–10 (2010).

12. Touchon, M. et al. Antibiotic resistance plasmids spread among natural isolates of Escherichia coli in spite of CRISPR elements. Microbiology+ 158, 2997–3004 (2012).

13. Gophna, U. et al. No evidence of inhibition of horizontal gene transfer by CRISPR–Cas on evolutionary timescales. The ISME Journal 9, 2021–2027 (2015).

14. Mangas, E. L. et al. Pangenome of Acinetobacter baumannii uncovers two groups of genomes, one of them with genes involved in CRISPR/Cas defence systems associated with the absence of plasmids and exclusive genes for biofilm formation. Microbial Genomics 5, 12 (2019).

15. Bikard, D., Hatoum-Aslan, A., Mucida, D. & Marraffini, L. A. CRISPR Interference Can Prevent Natural Transformation and Virulence Acquisition during In Vivo Bacterial Infection. Cell Host & Microbe 12, 177–186 (2012).

16. Zhang, Y. et al. Processing-Independent CRISPR RNAs Limit Natural Transformation in Neisseria meningitidis. Molecular Cell 50, 488–503 (2013).

17. Jorth, P., Whiteley, M. & Davies, J. E. An Evolutionary Link between Natural Transformation and CRISPR Adaptive Immunity. mBio 3, 53 (2012).

18. Touchon, M. et al. The genomic diversification of the whole Acinetobacter genus: origins, mechanisms, and consequences. Genome Biology and Evolution 6, 2866–2882 (2014).

19. Cooper, R. M. & Hasty, J. One-Day Construction of Multiplex Arrays to Harness Natural CRISPR-Cas Systems. ACS Synthetic Biology 9, 1129–1137 (2020).

20. Suárez, G. A. et al. Rapid and assured genetic engineering methods applied to Acinetobacter baylyi ADP1 genome streamlining. Nucleic Acids Research 48, 4585–4600 (2020).

21. Palmen, R., Vosman, B., Buijsman, P., Breek, C. K. D. & Hellingwerf, K. J. Physiological characterization of natural transformation in Acinetobacter calcoaceticus. Microbiology+ 139, 295–305 (1993).

22. Vink, J. N. A. et al. Direct Visualization of Native CRISPR Target Search in Live Bacteria Reveals Cascade DNA Surveillance Mechanism. Molecular Cell 77, 39–50.e10 (2020).

23. Lin, L., Ringel, P. D., Vettiger, A., Dürr, L. & Basler, M. DNA Uptake upon T6SS-Dependent Prey Cell Lysis Induces SOS Response and Reduces Fitness of Acinetobacter baylyi. Cell Reports 29, 1633–1644.e4 (2019).

24. Johnston, C., Martin, B., Polard, P. & Claverys, J.-P. Postreplication targeting of transformants by bacterial immune systems? Trends Microbiol 21, 516–521 (2013).

25. Liu, M. et al. The Clustered Regularly Interspaced Short Palindromic Repeat System and Argonaute: An Emerging Bacterial Immunity System for Defense Against Natural Transformation? Front Microbiol 11, 593301 (2020).

26. Zhang, Y. The CRISPR-Cas9 system in Neisseria spp. Pathog Dis 75, (2017).

27. Innamorati, K. A., Earl, J. P., Aggarwal, S. D., Ehrlich, G. D. & Hiller, N. L. The Bacterial Guide to Designing a Diversified Gene Portfolio. in The Pangenome, Diversity, Dynamics and Evolution of Genomes (eds. Tettellin, H. & Medini, D.) 51–87 (Springer, 2020). doi:10.1007/978-3-030-38281-0_3.

28. Bikard, D. et al. Exploiting CRISPR-Cas nucleases to produce sequence-specific antimicrobials. Nature Biotechnology 32, 1146–1150 (2014).

29. Bikard, D., Hatoum-Aslan, A., Mucida, D. & Marraffini, L. A. CRISPR Interference Can Prevent Natural Transformation and Virulence Acquisition during In^~^Vivo Bacterial Infection. Cell Host & Microbe 12, 177–186 (2012).

30. Stachler, A.-E. et al. High tolerance to self-targeting of the genome by the endogenous CRISPR-Cas system in an archaeon. Nucleic Acids Research 45, 5208–5216 (2017).

31. Caliando, B. J. & Voigt, C. A. Targeted DNA degradation using a CRISPR device stably carried in the host genome. Nat Commun 6, 6989 (2015).

32. Vercoe, R. B. et al. Cytotoxic chromosomal targeting by CRISPR/Cas systems can reshape bacterial genomes and expel or remodel pathogenicity islands. PLOS Genetics 9, e1003454 (2013).

33. McGinn, J. & Marraffini, L. A. CRISPR-Cas Systems Optimize Their Immune Response by Specifying the Site of Spacer Integration. Molecular Cell 64, 616–623 (2016).

34. Liao, C. et al. Modular one-pot assembly of CRISPR arrays enables library generation and reveals factors influencing crRNA biogenesis. Nature Communications 10, 2948 (2019).

35. Jones, D. L. et al. Kinetics of dCas9 target search in Escherichia coli. Science (New York, NY) 357, 1420–1424 (2017).

36. Høyland-Kroghsbo, N. M. et al. Temperature, by Controlling Growth Rate, Regulates CRISPR-Cas Activity in Pseudomonas aeruginosa. mBio 9, 1709 (2018).

37. Strotskaya, A. et al. The action of Escherichia coli CRISPR–Cas system on lytic bacteriophages with different lifestyles and development strategies. Nucleic Acids Research 28, gkx042 (2017).

38. Meeske, A. J., Nakandakari-Higa, S. & Marraffini, L. A. Cas13-induced cellular dormancy prevents the rise of CRISPR-resistant bacteriophage. Nature 570, 241–245 (2019).

39. Fukuyo, M., Sasaki, A. & Kobayashi, I. Success of a suicidal defense strategy against infection in a structured habitat. Sci Rep-uk 2, 238 (2012).

40. Murin, C. D., Segal, K., Bryksin, A. & Matsumura, I. Expression Vectors for Acinetobacter baylyi ADP1. Applied and Environmental Microbiology 78, 280–283 (2011).

