## Supplementary material for "CRISPR-Cas Inhibits Natural Transformation Through Altruistic Group Defense and Self-Sacrifice": Supplemental Figures.pdf

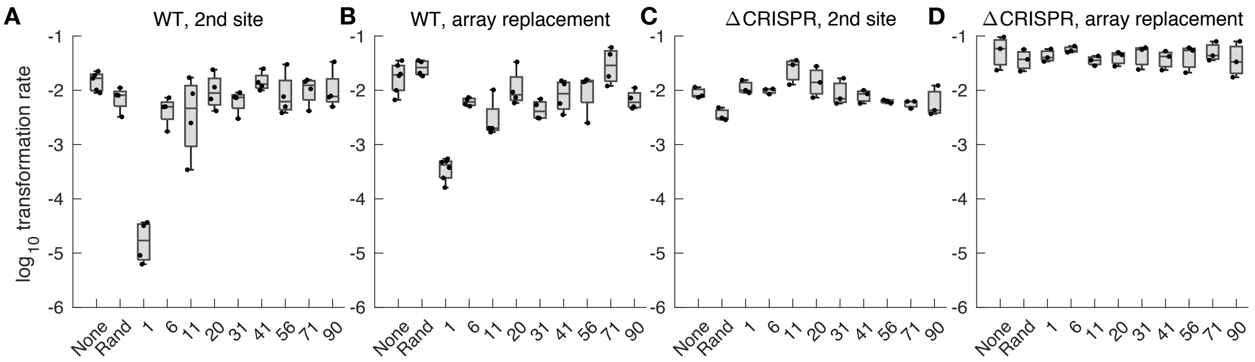


**Figure S1**: Natural transformation by DNA targeted by the endogenous CRISPR array. Transformation rates (transformants per recipient) are shown for donor DNA containing various targets matching spacers from the endogenous array, or a random spacer. A, C, wild type recipient cells; B, D, recipients lacking the CRISPR array. A,B, second-site donor DNA; C,D, donor DNA replaces the CRISPR array. Points indicate experimental replicates, which overlay standard box plots. Figure 1 shows the same data, but normalized by the no-target controls (the first box in each plot).

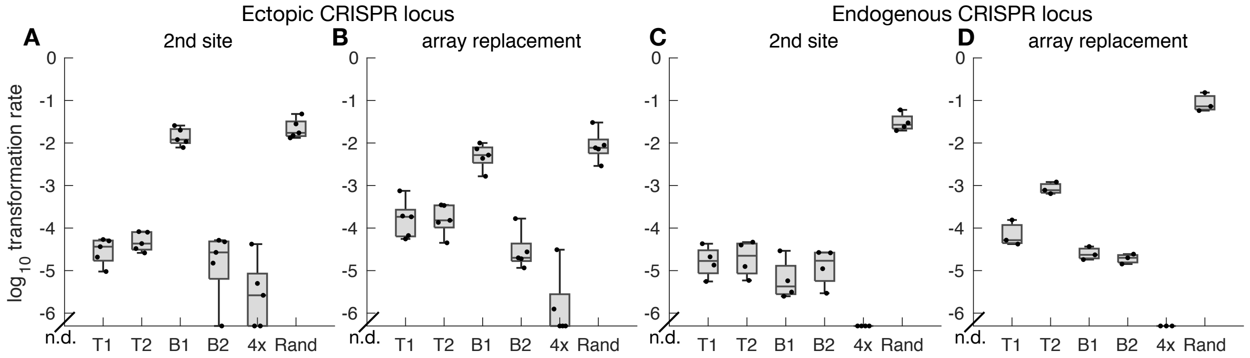


**Figure S2**: Natural transformation by DNA targeted by introduced CRISPR spacers. CRISPR spacers were introduced into an ectopic locus (A, B) or used to replace the endogenous locus (C, D). Donor DNA either inserted into a separate genomic site (A, C) or replaced the introduced spacers (B, D). Data points indicate experimental replicates, n=5 for A and B, n=4 for C, n=3 for D. n.d. = none detected.

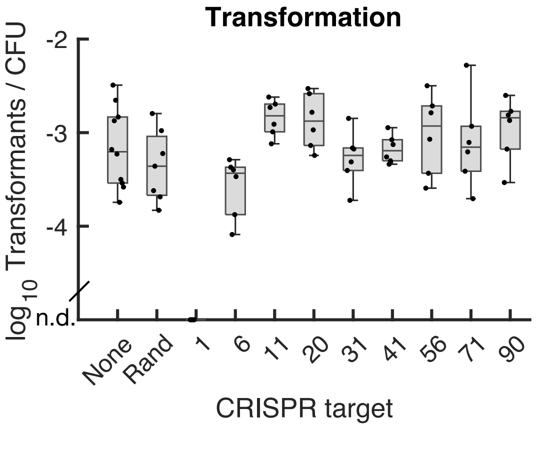


**Figure S3**: Transformation rate at the first time point of the pulse-chase experiments, expressed as transformants per recipient. This is the same data as in Figure 3, but not normalized by the no-target controls (here, CRISPR target "none"). Data below detection is shown along the x-axis, denoted n.d. for none detected.

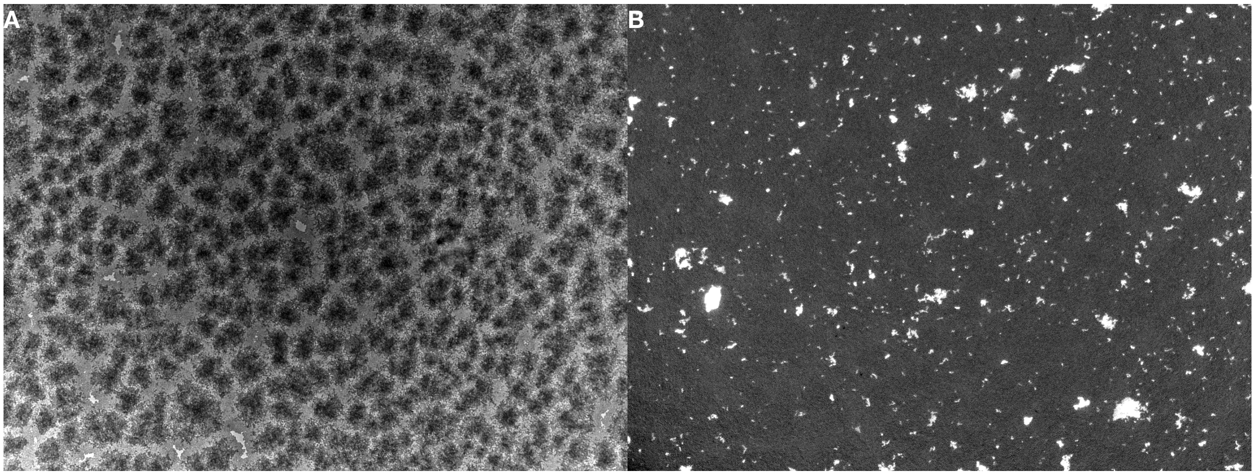


**Figure S4:** Transformation by a small transferring region on agar. Recipient cells containing a GFP gene broken by adjacent stop codons were incubated with ~3 kb donor DNA to repair the stop codons. A) transmitted light image of a filled field of view after 33 hours of growth; B) GFP channel.

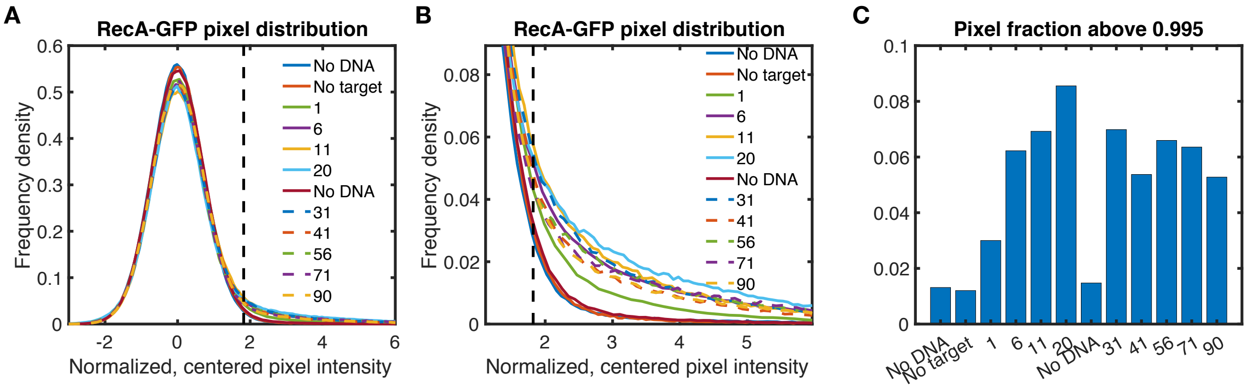


**Figure S5:** Quantifying DNA damage in images of *$P_{recA}$-GFP* cells. A) A representative set of background-corrected GFP pixel intensity histograms were normalized and centered, such that the lower half of each distribution fit a Gaussian curve $y = ae^{-x^2}$, with *a* chosen to normalize the entire histogram to area 1. B) A closer view of the tails from the curves in (A). A threshold was set at the 99.5 percentile of a normal curve (vertical dashed black line), and the tails beyond that threshold were considered to be GFP-positive. C) Fraction of GFP-positive pixels (area under the curves beyond the threshold) for the histograms in (A).

**Figures S6-14:** DNA damage reporter cells incubated under agar with the indicated donor DNA. Each image montage includes 6 experiments run in parallel on the same dish.

**Figures S15-S23:** Detected GFP-positive pixels for the corresponding GFP images in Figures S6-S14, respectively. Areas with no cells are outlined in red, and GFP-positive pixels are outlined in green.
