## Supplementary figures and images for "CRISPR-Cas Inhibits Natural Transformation Through Altruistic Group Defense and Self-Sacrifice"

### Figure S6.png

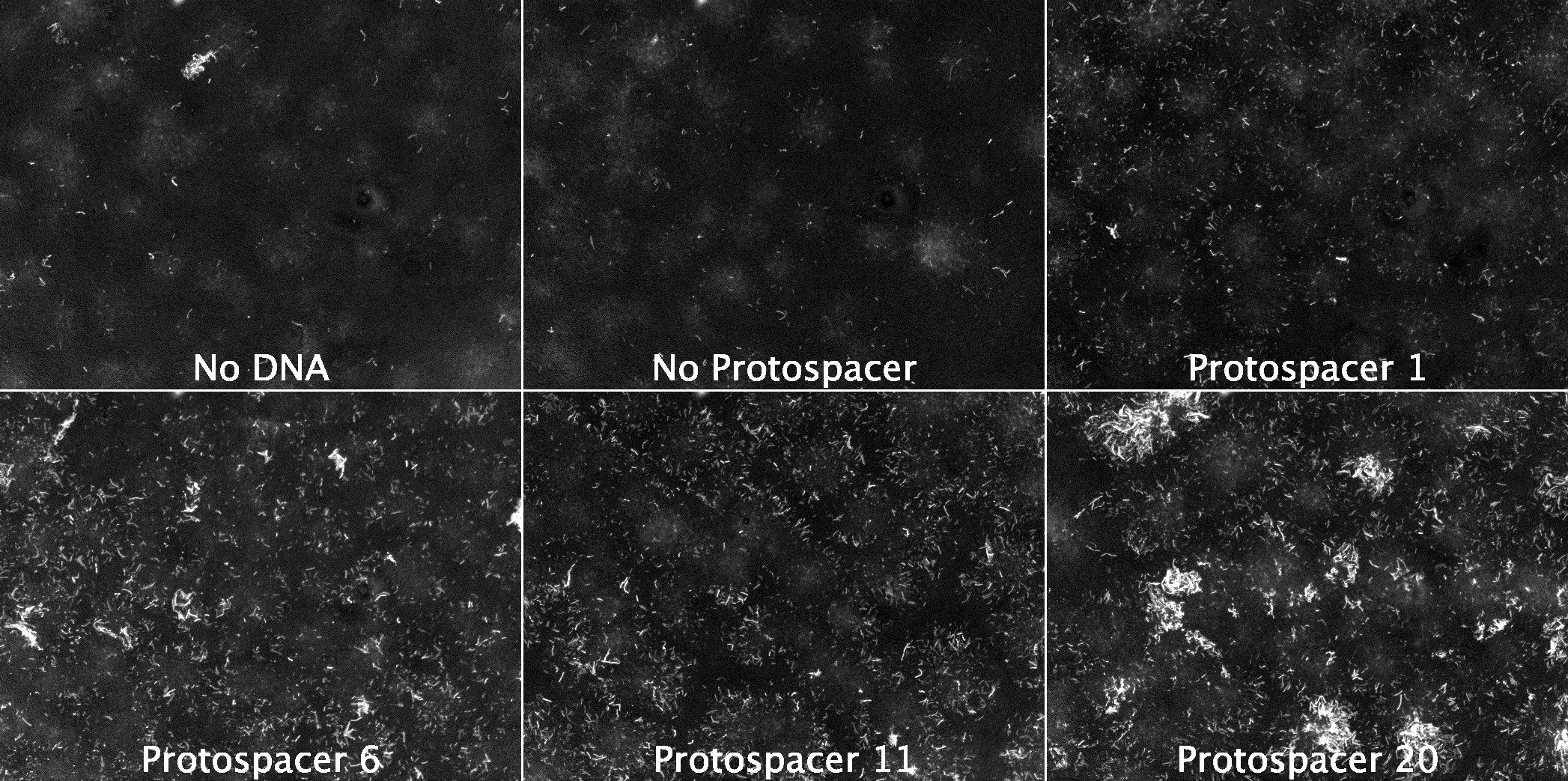

### Figure S7.png

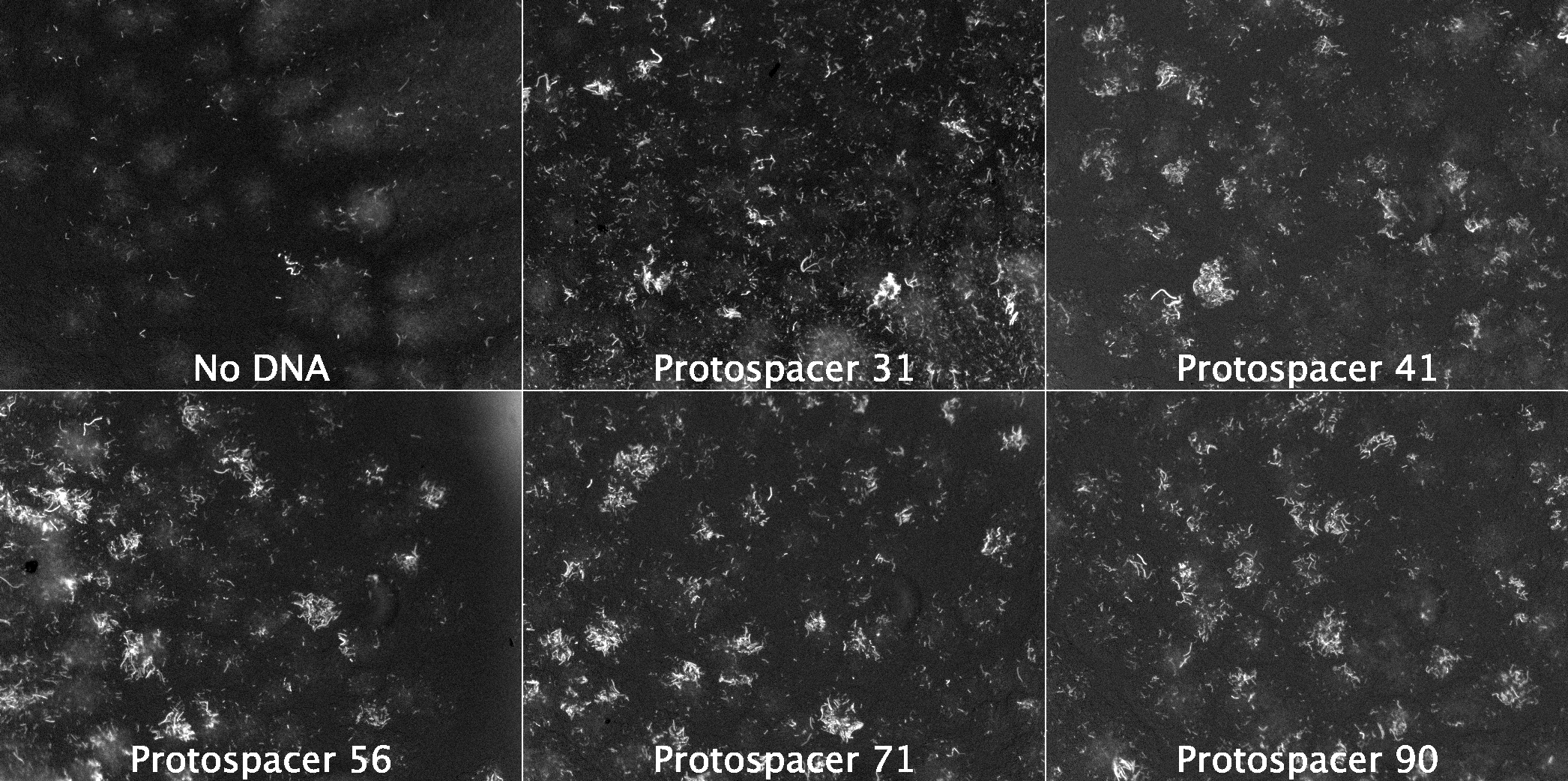

### Figure S8.png

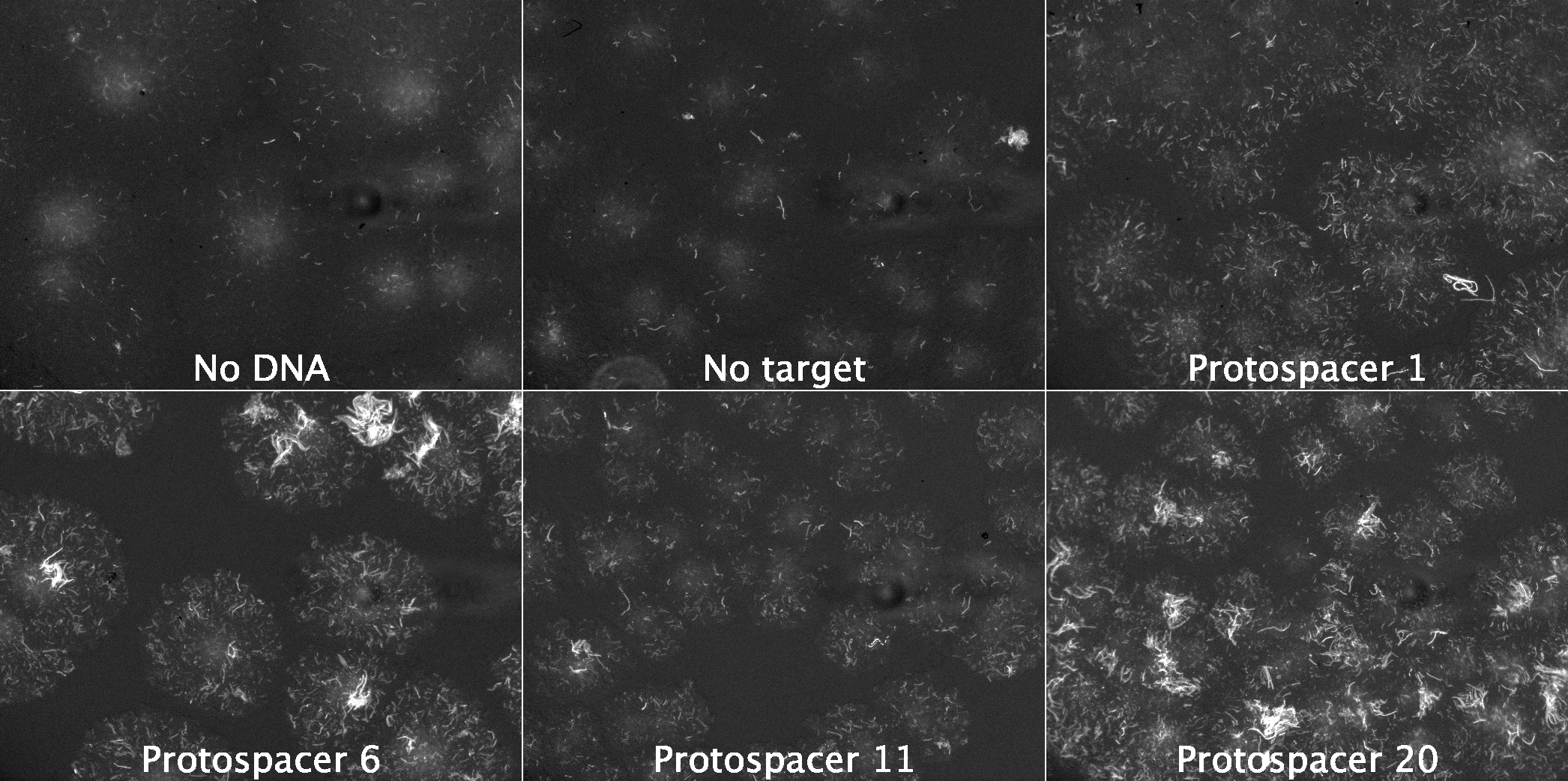

### Figure S9.png

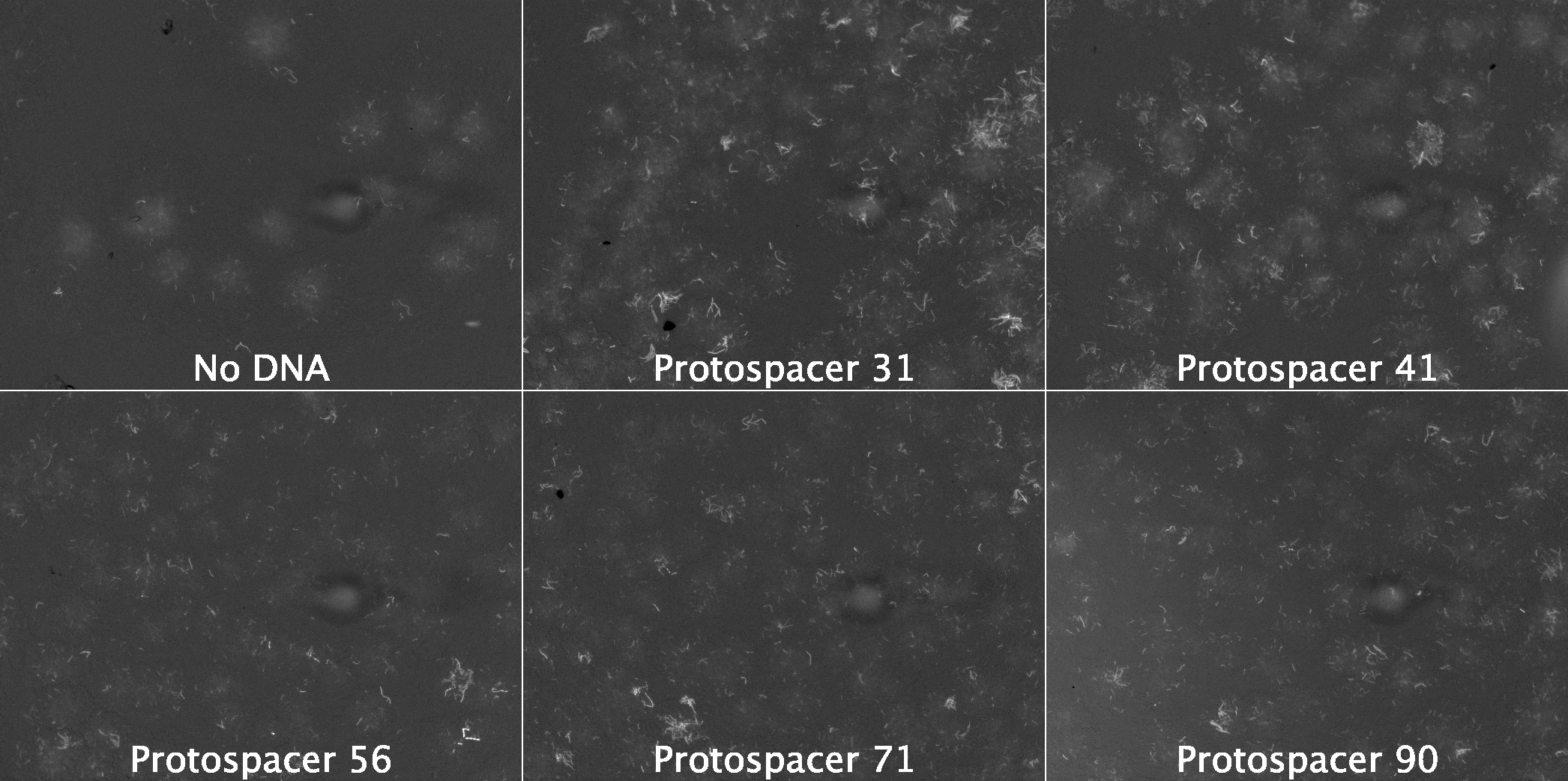

### Figure S10.png

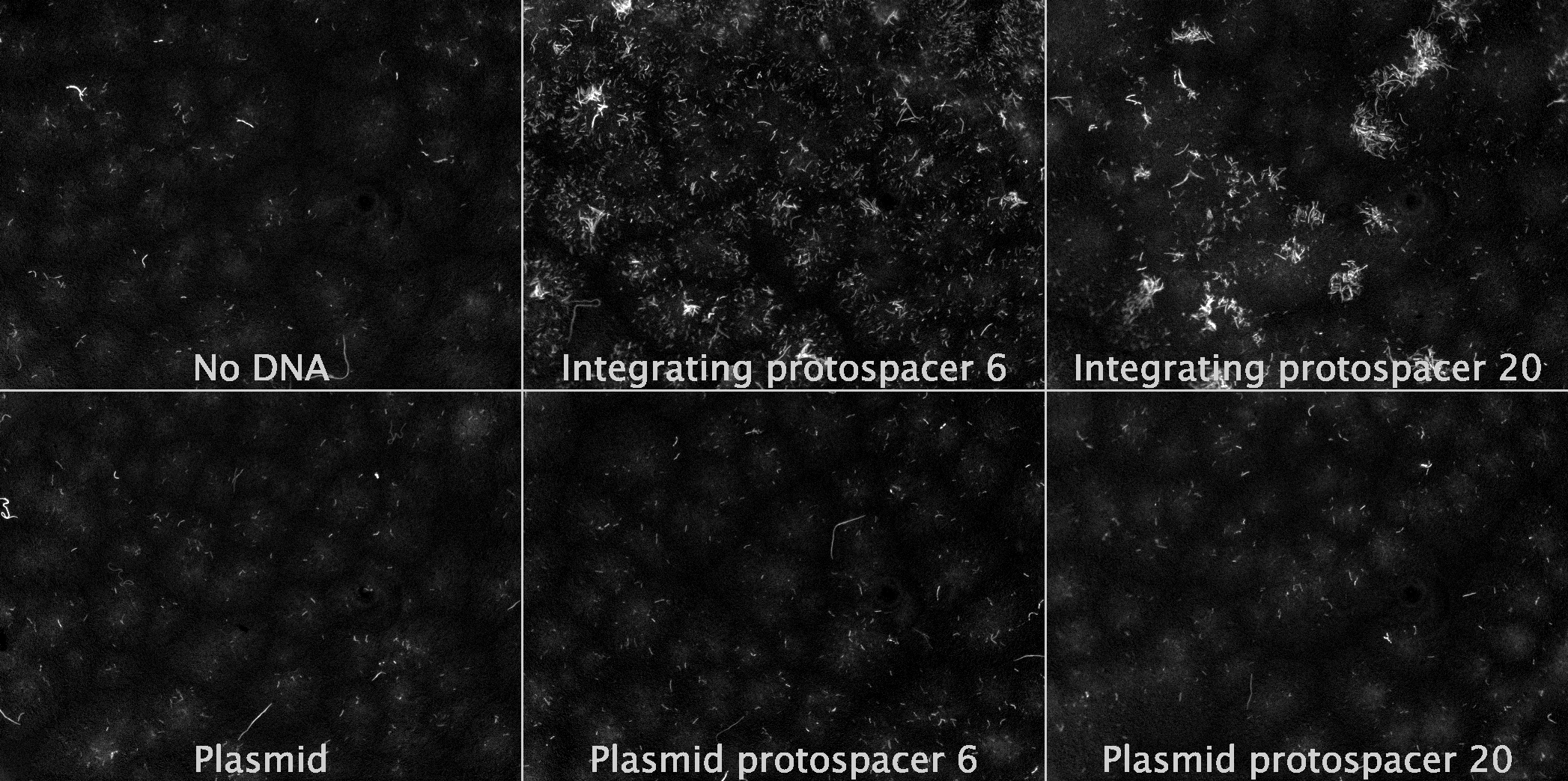

### Figure S11.jpg

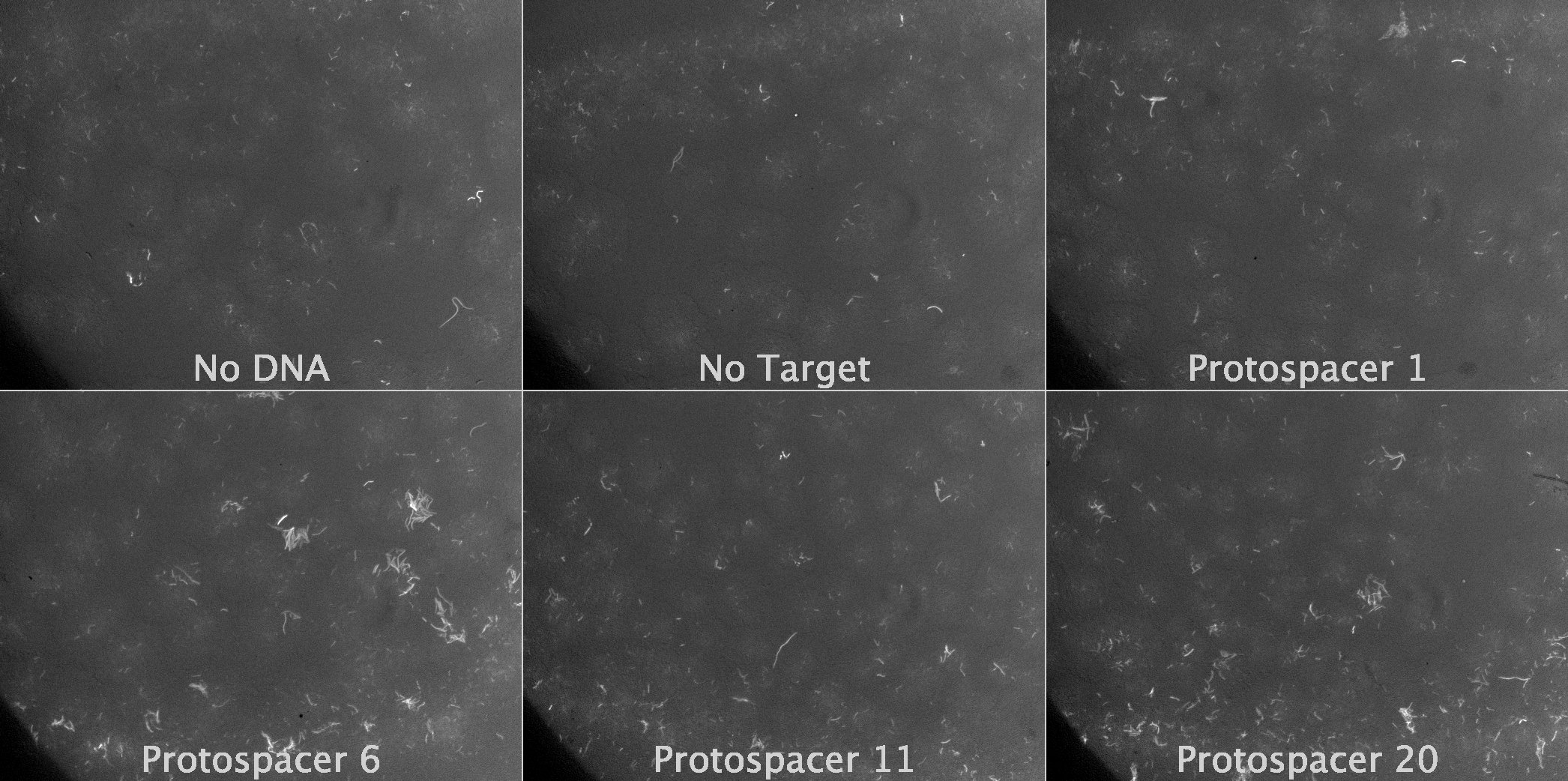

### Figure S12.jpg

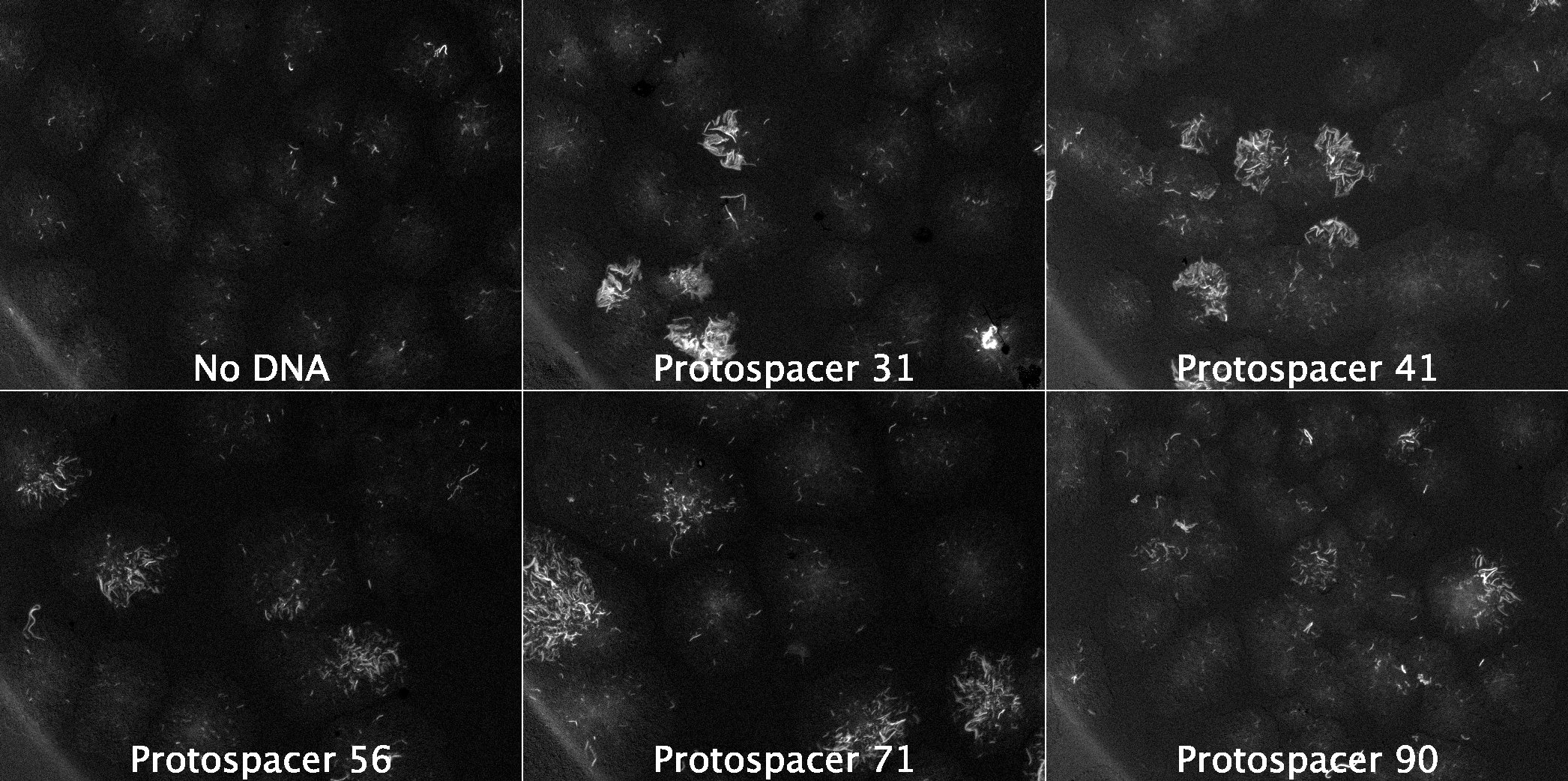

### Figure S13.jpg

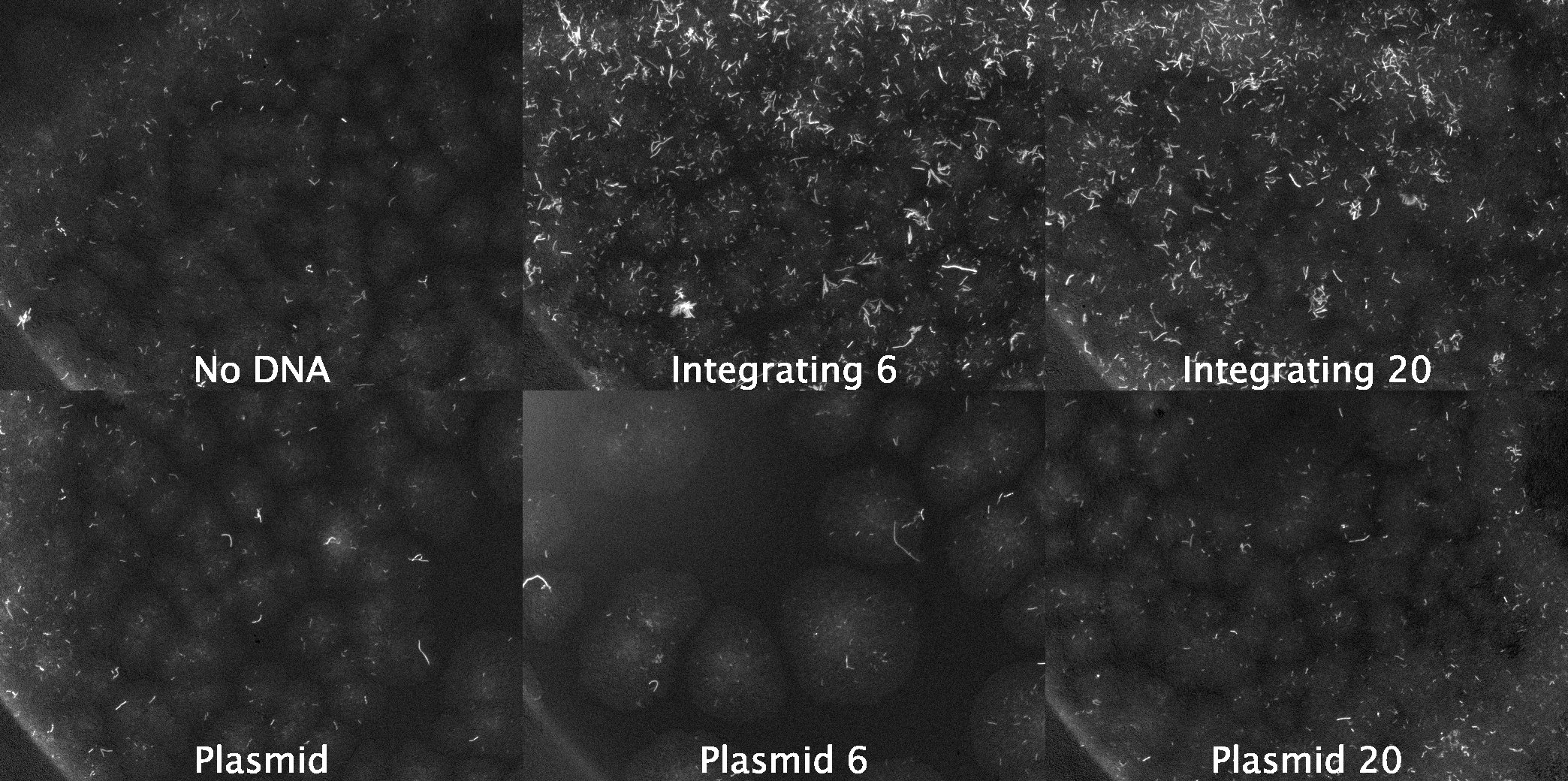

### Figure S14.png

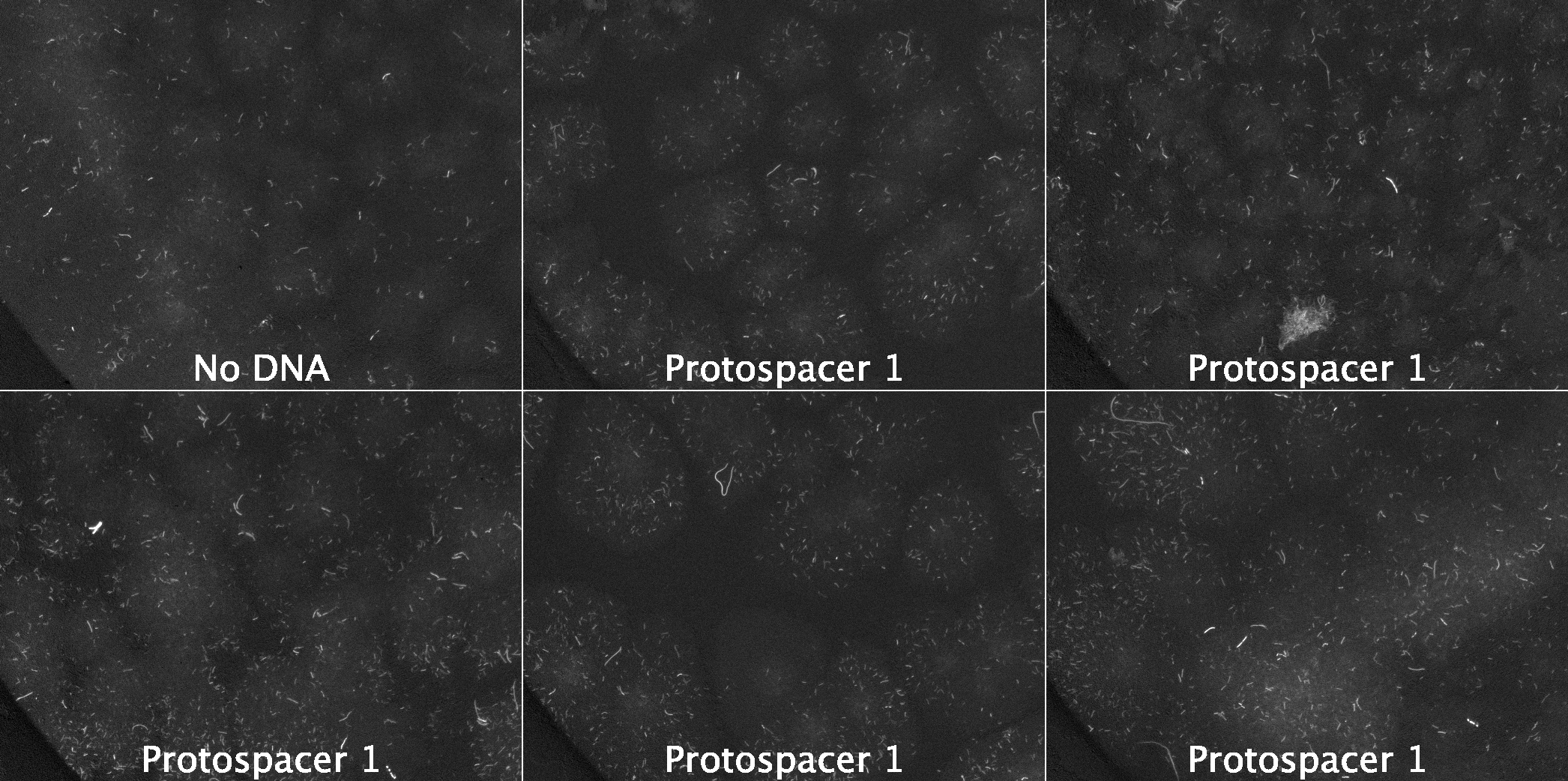

### Figure S15.png

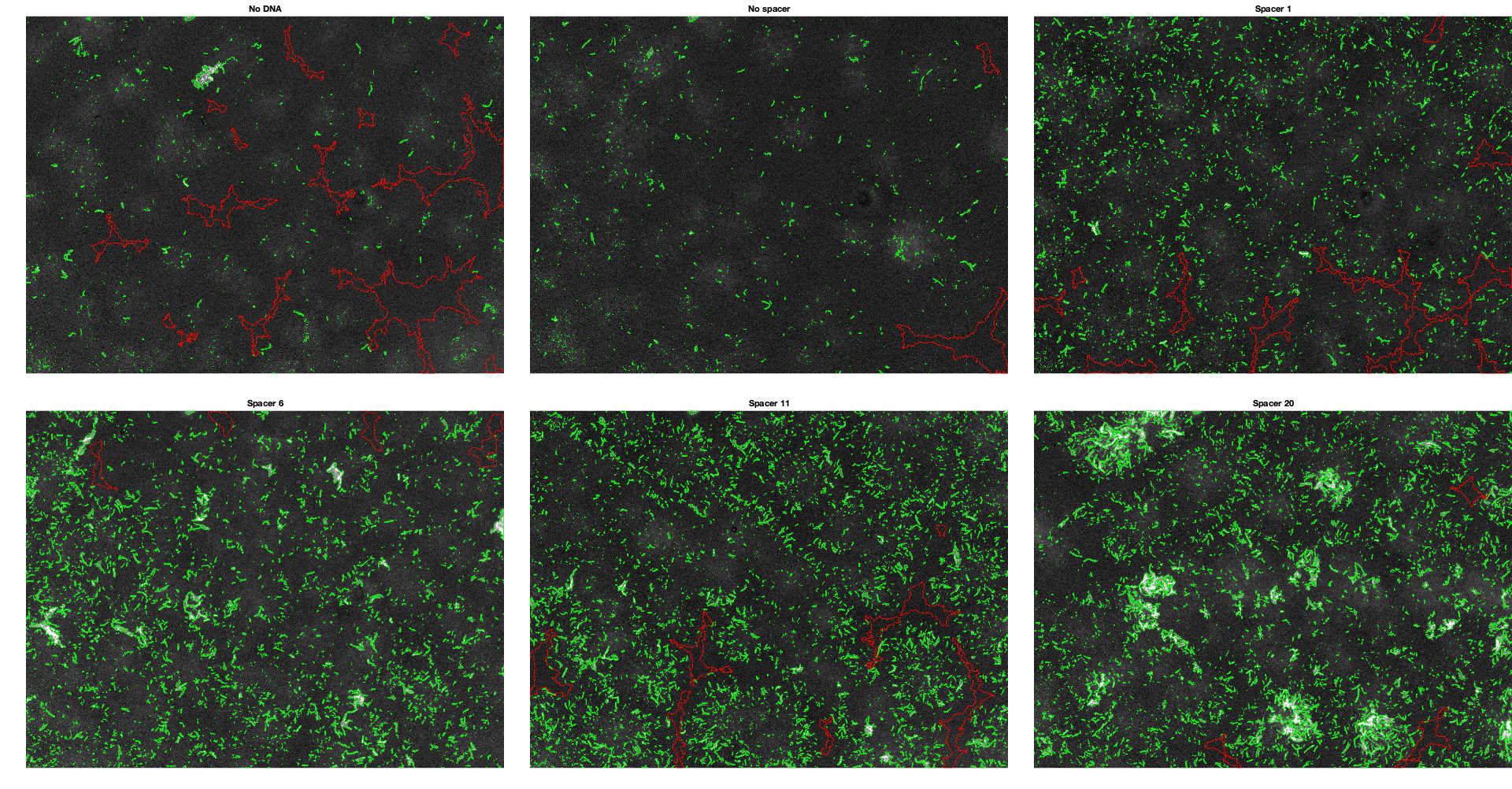

### Figure S16.png

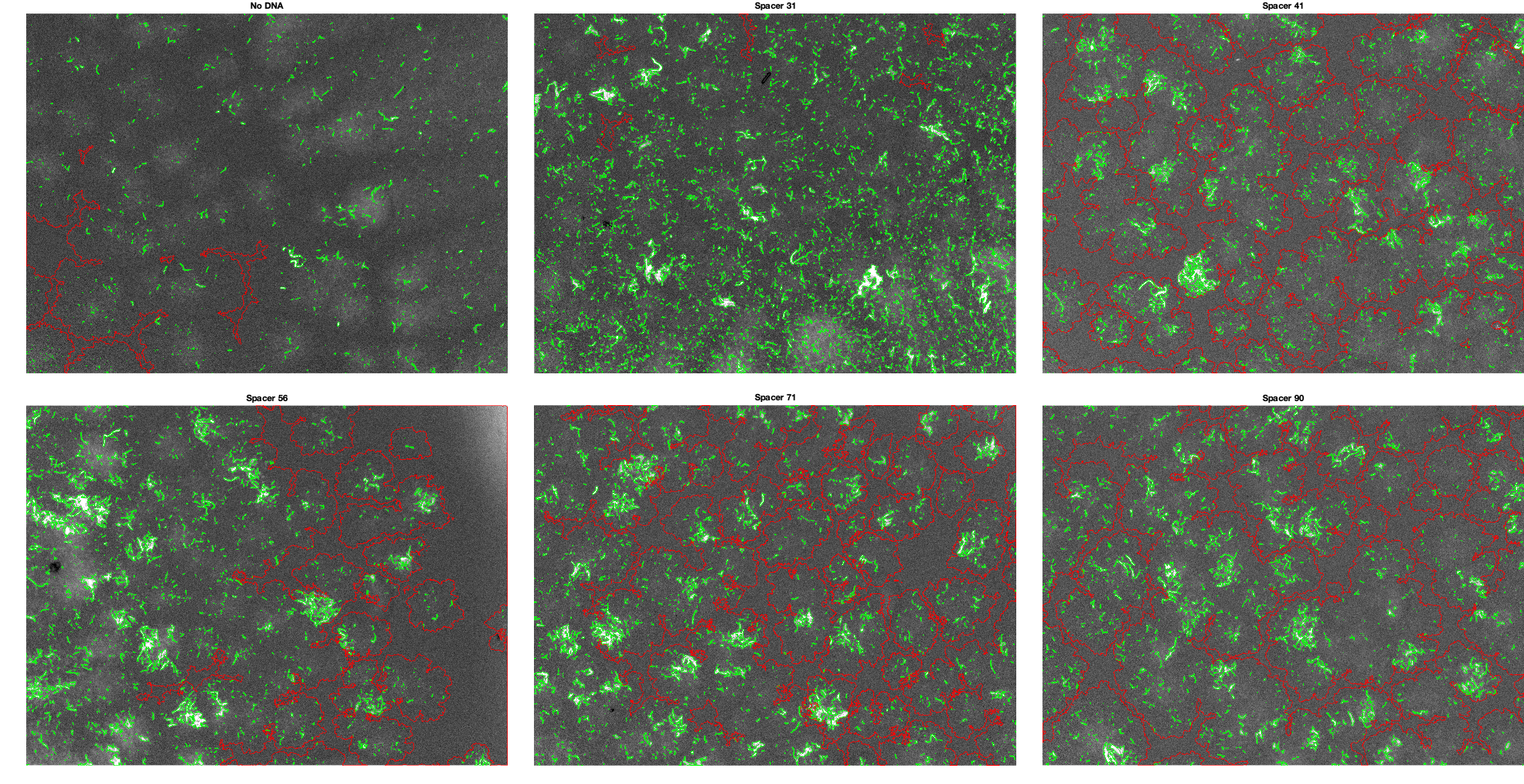

### Figure S17.png

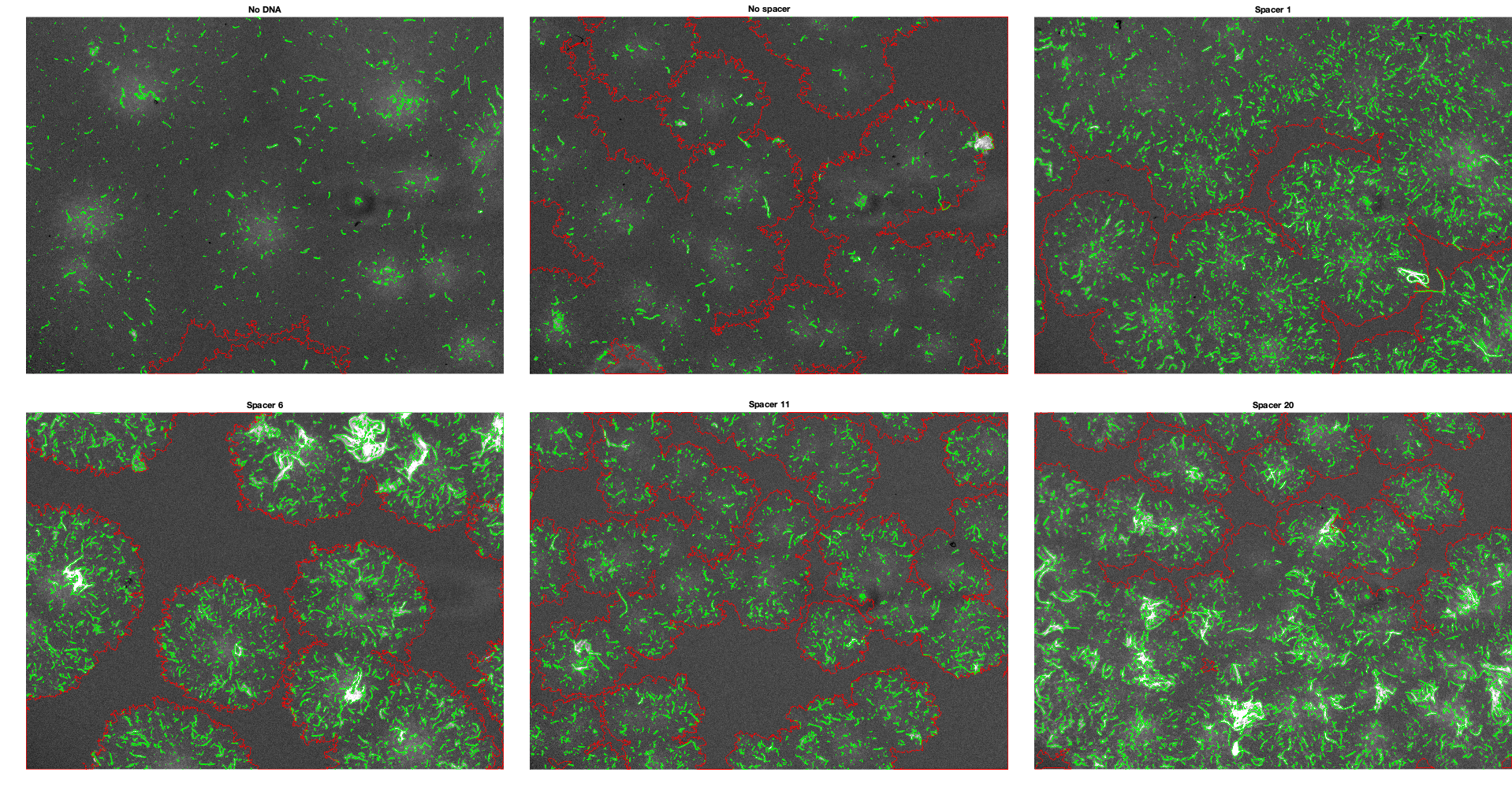

### Figure S18.png

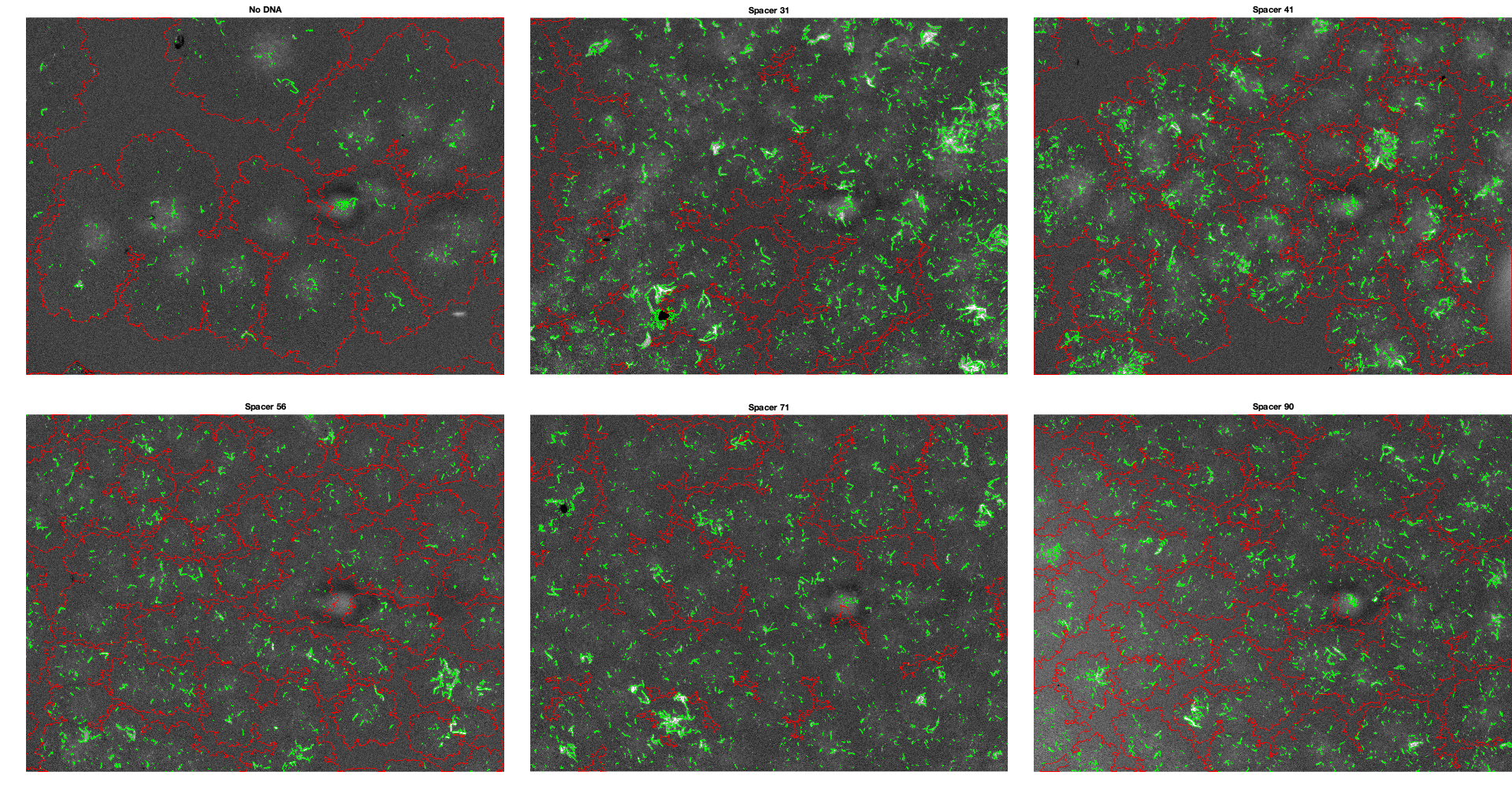

### Figure S19.png

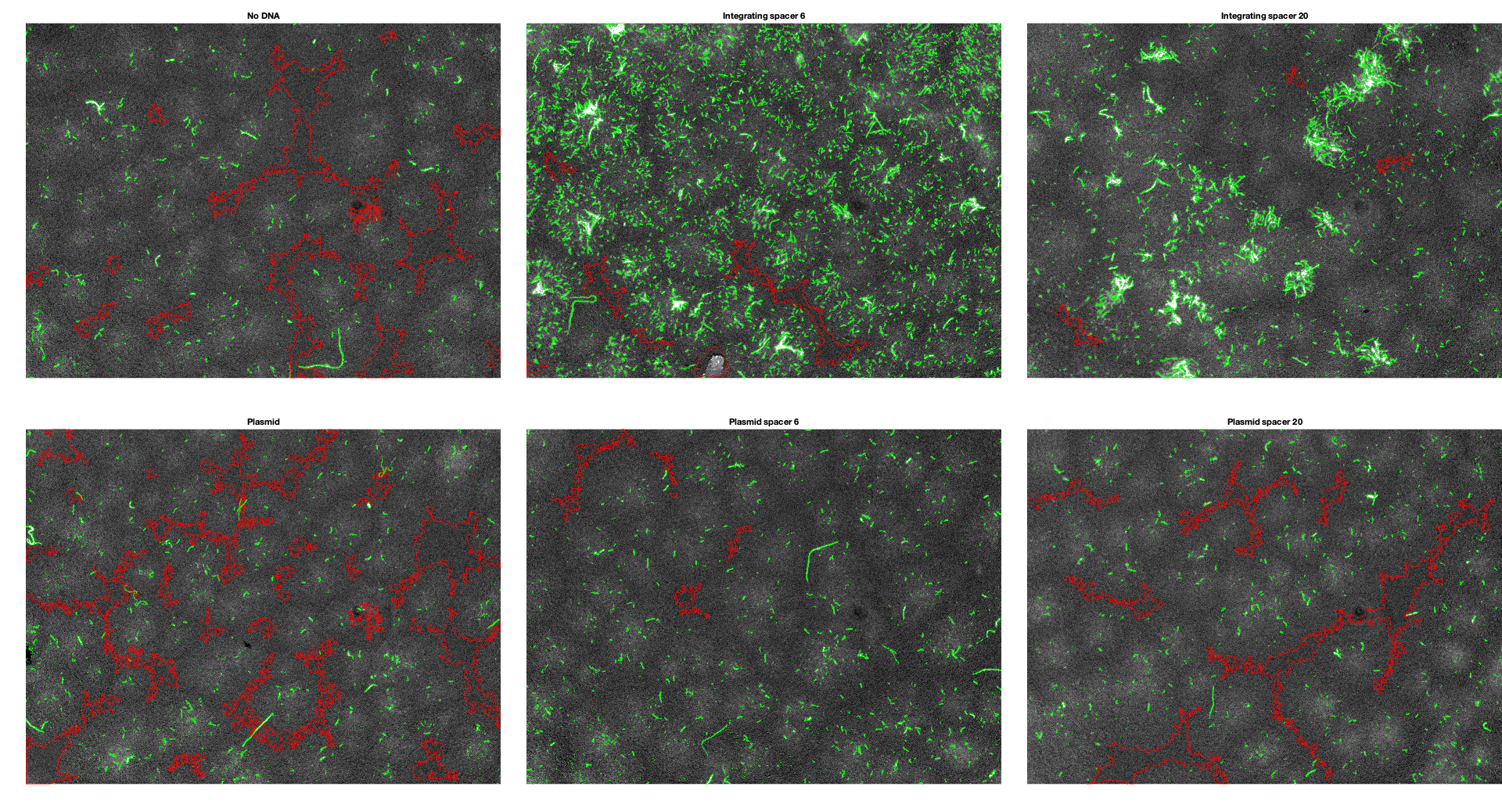

### Figure S20.png

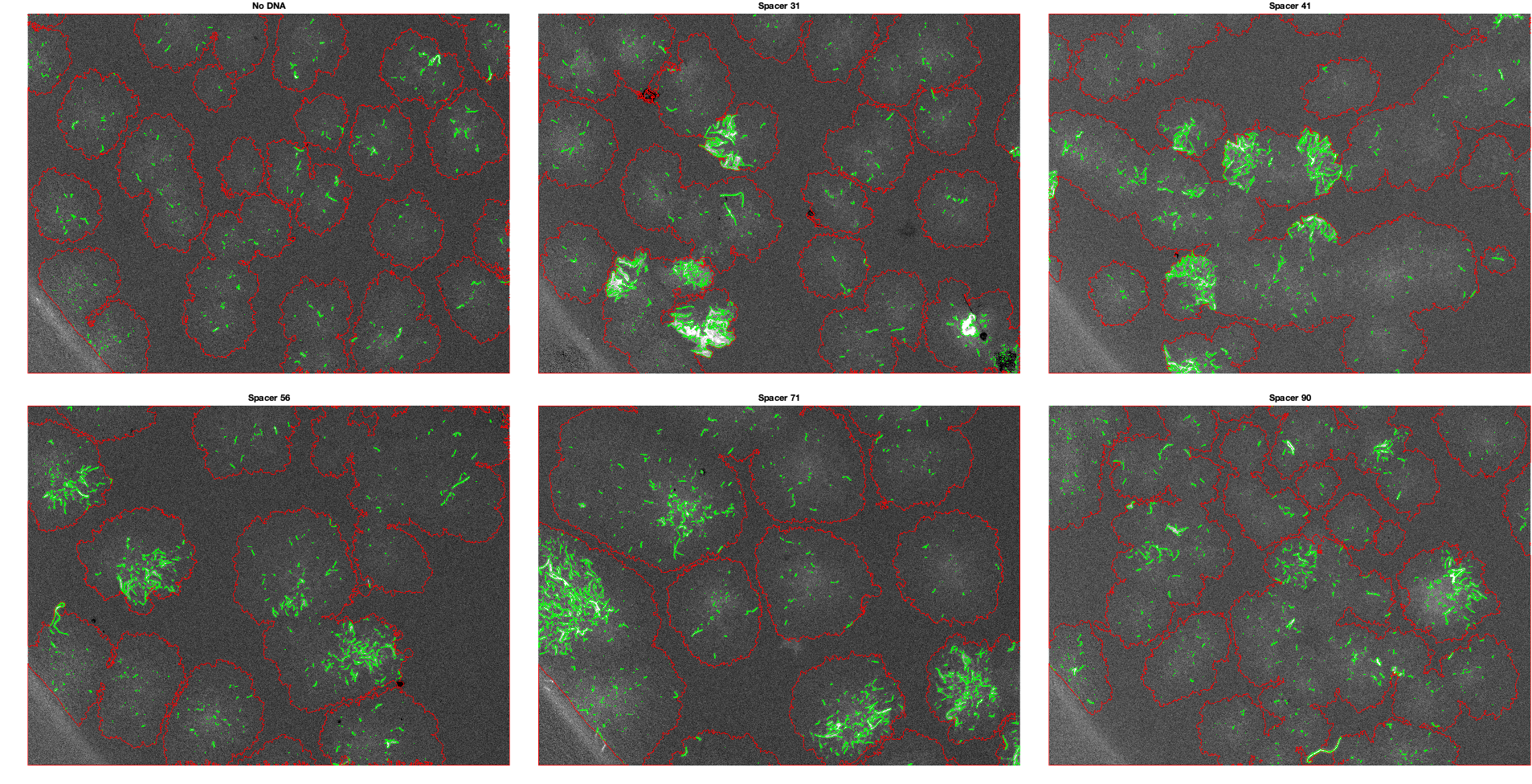

### Figure S21.png

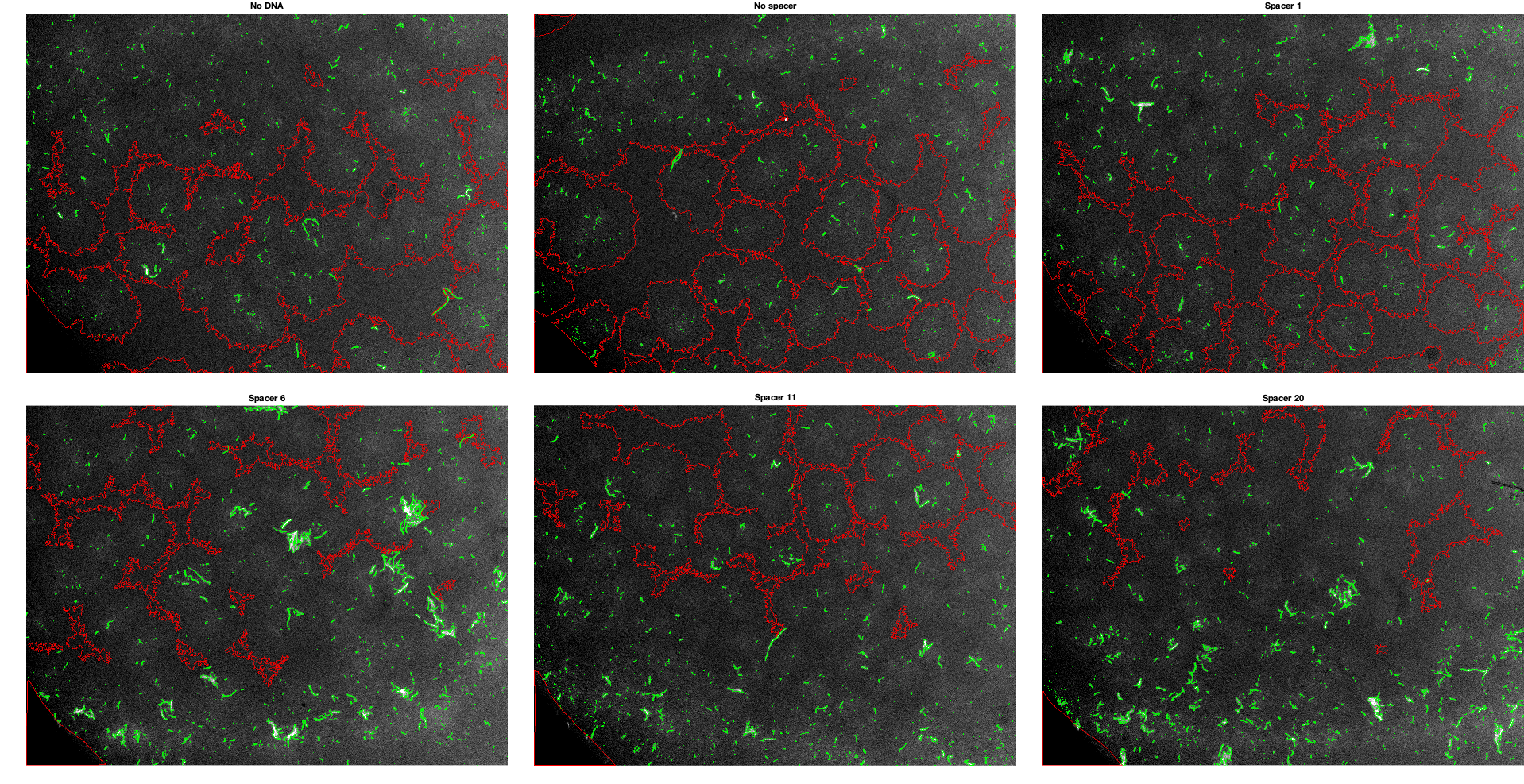

### Figure S22.png

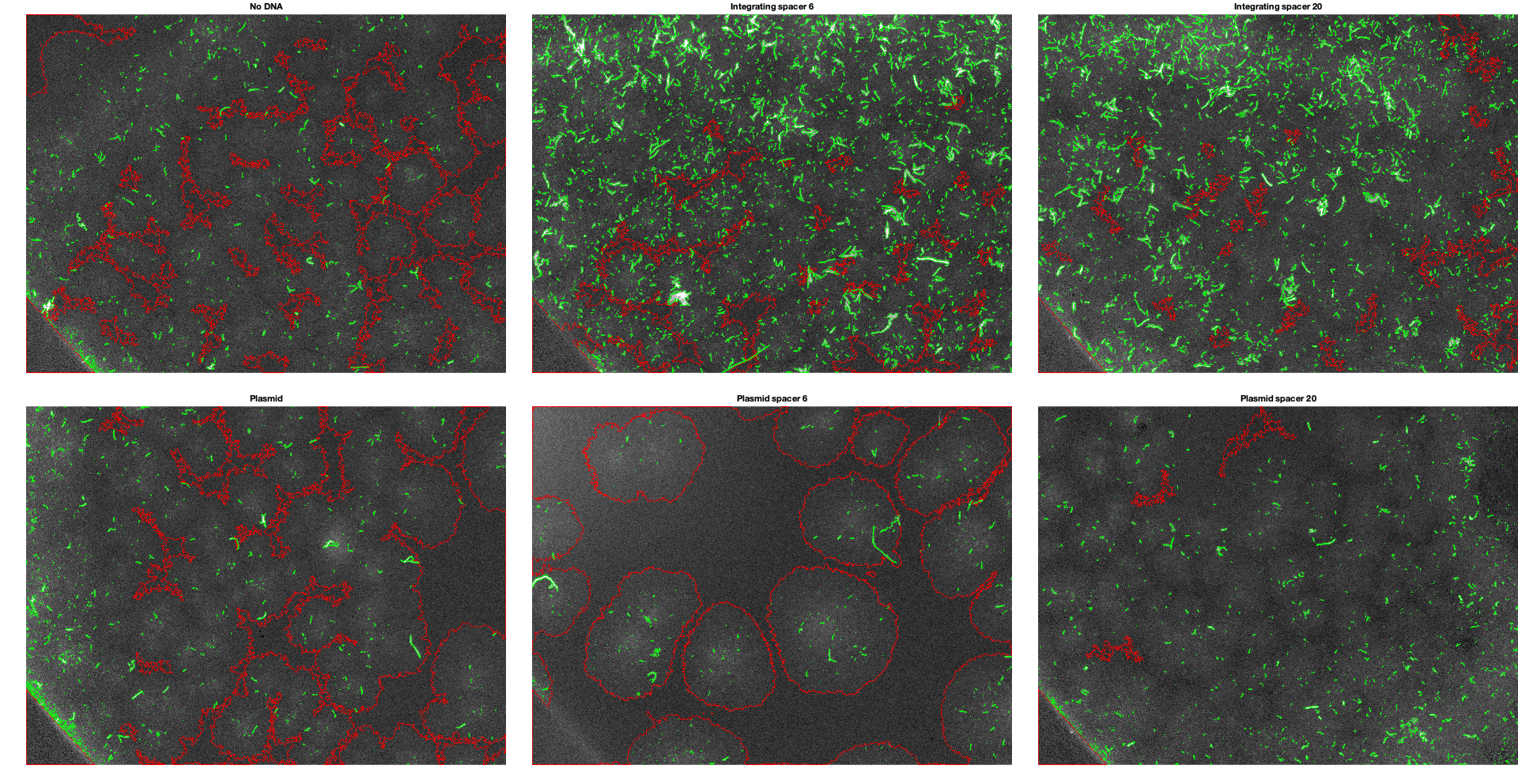

### Figure S23.png

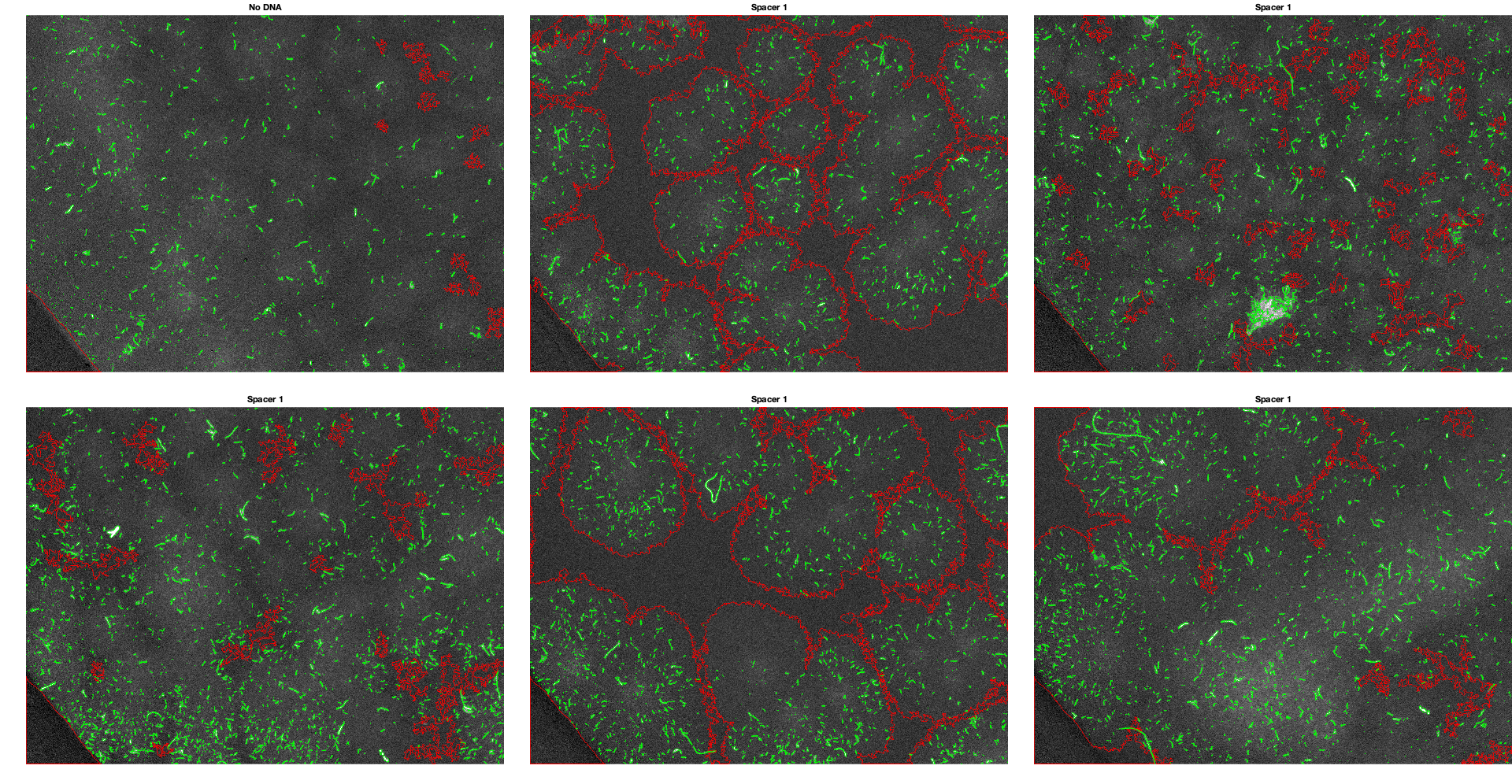
